# A Developmental Atlas of the *Drosophila* Nerve Cord Uncovers a Global Temporal Code for Neuronal Identity

**DOI:** 10.1101/2025.07.16.664682

**Authors:** Sebastian Cachero, Myrto Mitletton, Isabella R. Beckett, Elizabeth C. Marin, Laia Serratosa Capdevila, Marina Gkantia, Jelly H.M. Soffers, Haluk Lacin, Gregory S.X.E. Jefferis, Erika Donà

## Abstract

The assembly of functional neural circuits relies on the generation of diverse neural types with precise molecular identity and connectivity. Unlocking general principles of neuronal specification and wiring across the nervous system requires a systematic and high-resolution characterisation of its diversity, recently enabled by advances in single-cell transcriptomics and connectomics. However, linking the molecular identity of neurons to circuit architecture remains a key challenge. Here, we present a high-resolution developmental transcriptional atlas for the *Drosophila melanogaster* nerve cord, the central hub for sensory-motor circuits. With an unprecedented 38x aggregate coverage relative to its reference connectome^1,2^, our atlas captures extensive molecular diversity and enables robust alignment to the adult connectome. We identified three developmental principles underlying neuronal diversity in the nerve cord. First, timing of neurogenesis shapes diversification of molecular identity: embryonic-born neurons diverge faster than larval-born neurons, as also observed in the adult connectome. Second, 17 transcription factors common to neurons from all lineages provide a global molecular identity code for birth order. Lastly, by mapping sex-specific transcriptional profiles to the connectome, we uncovered female-specific apoptosis and transcriptional divergence as key global drivers of sex-specification. By revealing key organisational axes of molecular identity, this atlas opens new avenues to dissect the molecular mechanisms underpinning the development and evolution of neural circuits.

## Introduction

The brain’s ability to process information and produce adaptive behaviors relies on its cellular composition and circuit organisation. Throughout development, neurons adopt specific molecular identities, which define their physiological function, and guide their integration into precise circuit architectures. While spatial and temporal patterning mechanisms coordinating neuronal fate are starting to be elucidated^3–5^, it is increasingly clear that many jigsaw pieces are still missing to account for the vast cellular complexity of the brain. To gain a comprehensive understanding of how neurons are specified across the nervous system we need a systematic and high-resolution characterisation of this diversity. Over the past decade, this need has driven two types of large-scale atlasing efforts: single-cell transcriptomics to define the molecular identities^6–11^, and connectomics to reveal anatomical and synaptic relationships of neurons^1,12–15^.

In both approaches the dimensionality of the large datasets can be reduced by grouping individual neurons into cell types which represent the functional units of neuronal circuits^16^. Crucially, a mature taxonomy of cell types requires refinement by integration across modalities^17,18^. In vertebrates, recent mouse and human brain atlases^7–10^ have identified more than 5,000 and 3,000 transcriptional types respectively^17,18^. However, the establishment of a comprehensive correspondence between transcriptional and anatomical types is hampered by the lack of reference connectomes. The situation is different for the highly stereotyped and numerically more modest brain of Drosophila melanogaster. Here, the systematic analysis and annotation of the combined datasets of the central nervous system, the largest connectomes yet completed, revealed over 11,000 neuronal types across 160,000 neurons^2,19,20^. This complete description of the texture of a complex nervous system provides an exciting opportunity to better understand the development of neural circuits.

However, comprehensive cross-identification of anatomical and molecular cell types remains challenging. The most complete matching to date has been obtained in sensory systems: for olfactory projection neurons^21^ and in the optic lobe^22–26^, where ∼800 repeated ommatidia result in hundreds of copies for a relatively small number of cell types. This unusually high abundance greatly facilitates the recovery of transcriptional clusters. Indeed, the latest version of the optic lobe atlas has identified 233 transcriptional types and cross-matched 113 to the connectome^23^, even though 438 types intrinsic to the optic lobe have been annotated^27^. Notably, the rarest of these cell types identified in transcriptional data is present in 16 copies per hemisphere; just 29 out of the 10,910 cell types in the rest of the CNS are present in such large numbers. With most CNS cell types represented by ∼2 neurons per hemisphere, transcriptional coverage becomes a limiting factor: empirical observations and simulation suggest a minimum of 20-30 cells are needed to identify a cell type in scRNAseq datasets^17,28^. As a consequence, low-coverage atlases are inherently unsuited to resolve the full connectomic diversity, explaining why large-scale atlases outside the optic lobe^29–31^ reported ten times fewer neuronal types^2,17,19^, and only matched a minority to the connectome.

High coverage alone, however, might not be sufficient to resolve connectomic types, as work in the olfactory and visual system has shown that transcriptional diversity peaks during development and decreases in adults^5,21,25,32^, making developmental atlases better suited to capture the diversity in the connectome.

Stimulated by these considerations, we have obtained a developmental transcriptional atlas of the *Drosophila melanogaster* VNC, which houses the sensory-motor circuits that form the core interface between the nervous system and the body. The atlas’s deep coverage (38x aggregate coverage) enabled high-resolution mapping of neuronal diversity as well as emergent organisational features of molecular identity; comprehensive annotation allowed most cells to be linked to the connectome. Together, these properties of the atlas enabled the identification of a global temporal coordinate system: a sequence of 17 transcription factors (TFs) that define a persistent lineage-agnostic molecular correlate of neuronal birth order. Comparison of male and female data identified apoptosis and transcriptional divergence as key developmental mechanisms of sex differences and also enabled precise matching with connectomic cell types. This atlas provides a resource and intellectual framework for uncovering new ontogenetic principles of circuit assembly.

## Results

### Developmental transcriptional atlas of the fly nerve cord

To reveal the transcriptional diversity of the 16,000 intrinsic neurons of the VNC we built an atlas using single cell RNA sequencing (scRNAseq) (Extended Data Fig. 1, Methods). After quality control we obtained 459,091 cells from four developmental stages (6, 24, 36 and 48h after puparium formation), spanning the period when neuronal morphology and connectivity are established^33^ (Fig. 1a). The atlas includes a balanced number of male and female cells, making it suitable for studying the development and function of sexually dimorphic circuits (Fig. 1b).

Dimensionality reduction produced a complex embedding in UMAP space, in line with the large number of cell types present in the VNC (Fig. 1b). We started annotating the atlas by separating neurons from glia, using known markers (e.g. *repo* and *wrapper* for glia and *elav* for neurons) (Fig. 1b, Methods). The ratio of glia to neurons moderately but significantly increased from 6 to 48h (Fig. 1b) matching reports of gliogenesis occurring in early pupal stages^34^, and the presence of actively dividing glial subpopulations in our atlas (Extended Data Fig. 2). Hereafter, we treated these cell populations independently to maximise cell type separation. While we focused primarily on neurons, we annotated six main glial types based on previously reported markers (Extended Data Fig. 2, Methods).

In the neuronal dataset, one of the main drivers of transcriptional diversity is maturation, which results in layered organisation of neurons in UMAP space according to the developmental stage at which they were sampled (Fig. 1c). This pattern is mirrored by graded expression of nSyb, a synaptic protein commonly associated with neuronal maturation (Fig. 1ci-ii). While this structure reflects molecular dynamics of cell-specific maturation, it hinders the identification of matching cell types across development.

**Figure 1.**
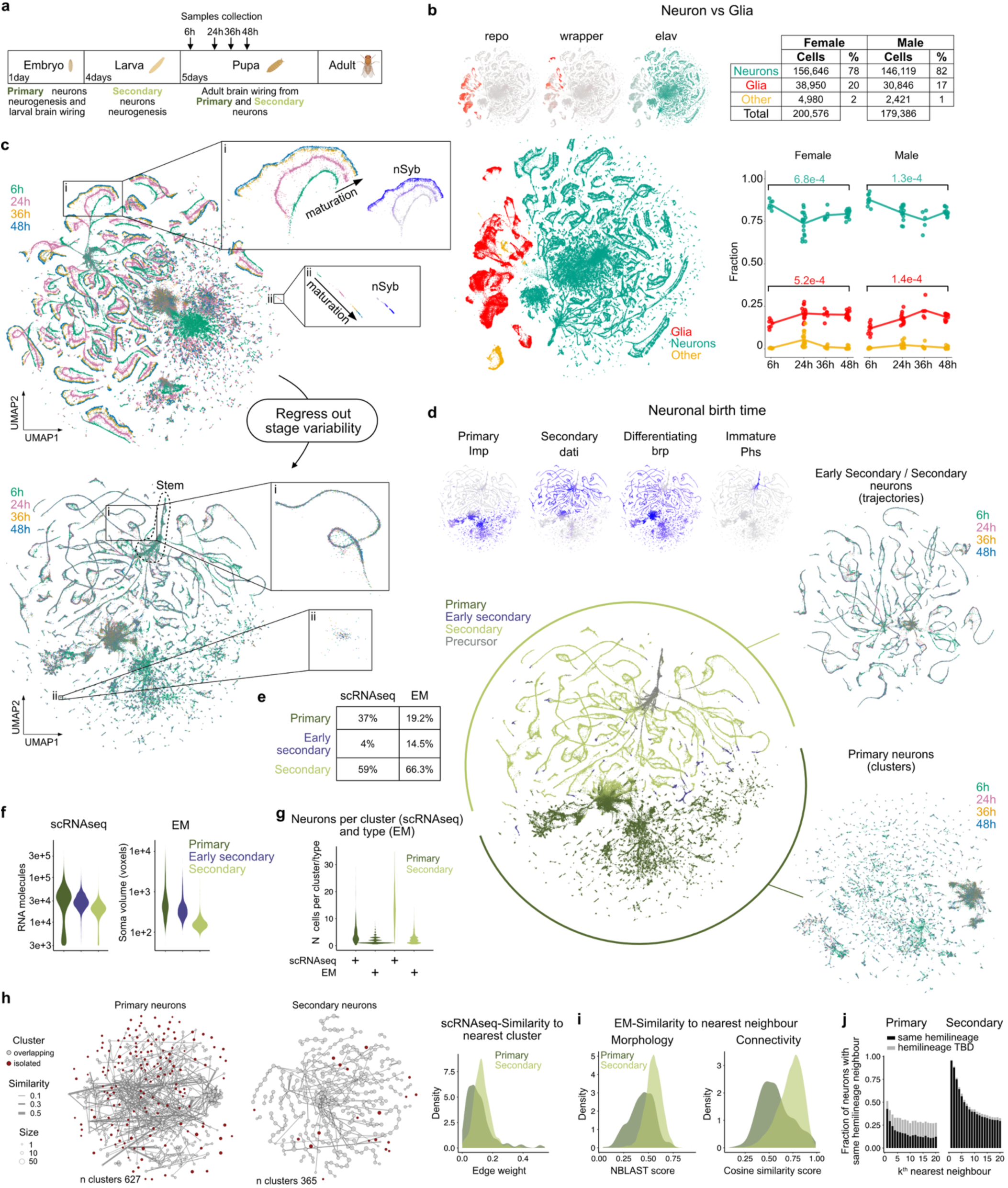
A developmental atlas of the *Drosophila* VNC uncovers molecular differences between primary and secondary neuronal types mirrored by the connectome. **a**, Timeline of *Drosophila* brain development highlighting two neurogenetic periods (embryonic and larval). Arrows: stages sampled for making the atlas. **b**, Top: UMAP plots showing the expression of glial and neuronal marker genes. Bottom: atlas annotation by glia, neurons and other (cells of mesenchymal origin). Table showing the number of glia, neurons and other cell types after demultiplexing and quality control. The graph shows the fraction of each population in the sampled stages split by sex. Each point corresponds to a single animal, brackets indicate statistical significance determined by t-test. **c**, Top: UMAP embedding for the neurons in the atlas coloured by stage. Bottom: same as top after regressing out maturation-driven variability using the 24h data as reference. Insets highlight two subsets of cells (i and ii respectively) in either embedding. nSyb expression is shown in the non-integrated embedding, showing that expression increases with stage. The “stem” region is indicated by a dotted line. **d**, Top: UMAP plots showing expression of markers of primary, secondary, differentiating neurons and precursor cells. Bottom: UMAP plot coloured by those categories. Plots on the right show UMAP embedding obtained by re-analysing primary and secondary neuronal subsets independently. **e**, Fractions of primary, secondary and early secondary neurons in the atlas and the connectome. **f**, Violin plots showing the number of RNA molecules (UMIs) per neuron in the atlas (left) and neuronal soma volume per neuron in the connectome (right) according to neurogenesis wave categories. **g,** Number of neurons per scRNAseq cluster after normalising for expected coverage (coverage without 6h stage is 28x) and per connectomic type (see Methods) in primary and secondary neurons. **h,** Constellation plots showing inter-cluster overlaps in primary and secondary neurons. Nodes: cluster centroids (size proportional to cluster size normalised by 28x coverage). Brown = isolated clusters after minimum weight filtering (threshold = 0.1). Edges: overlap between clusters, width is proportional to the summed fraction shared between the two clusters, in both directions, i.e. for a given A->B, the edge is the fraction of A nn in B + the reverse (see Methods). Density plot shows distribution of the strongest edge for each cluster (number of clusters, primary = 627, secondary = 365, Wilcoxon rank-sum test, p < 1.3e-14). **i**, Left: density distribution of the NBLAST morphology scores to the top-match for primary and secondary neurons from thoracic segments in the male connectome (primary: n = 1,386, median = 0.45; secondary: n = 9,919, median = 0.55). Right: density distribution of the cosine similarity connectivity score to the nearest neighbour for primary and secondary neurons from thoracic segments in the male connectome (primary: n = 1,356, median = 0.55; secondary: n = 9,726, median = 0.74). For both comparisons, secondary neurons show significantly higher similarity than primary neurons (Wilcoxon rank-sum test, p < 2.2e-16). j, Fraction of neurons with the same hemilineage as the k^th^ neighbour at increasing neighbor rank (k), with neighbors ordered by decreasing similarity. Bars represent the cumulative fraction of neurons with the same hemilineage (black), as well as those for which the hemilineage of the neighbour is undetermined (TBD).

We therefore regressed out maturation-related variability, enabling the integration of transcriptomes from the same cell types across different stages, effectively aligning neurons by identity rather than developmental stage (see Extended Data Fig. 9c). One exception is the “stem” region, which contains less differentiated cells from 6 and 24h samples that do not integrate with 36 and 48h (Fig. 1c). Our neuronal atlas contains 302,765 cells representing the 7,880 neurons per one lateral half of the VNC connectome, yielding an unprecedented 38x aggregate coverage (Extended Data Fig. 1a,f for per-stage coverage, Methods-Connectomics data analysis).

### Embryonic and larval neurogenesis result in distinct modes of neuronal diversification

A common feature of nervous system development is that neuronal populations have different temporal origins. One example of this in insects is the existence of two waves of neurogenesis. Primary neurons are born in the embryo where they build the larval brain, but are then remodelled through pruning and regrowth of arbours during metamorphosis. Later, a subset of the same pool of stem cells produces secondary neurons during larval stages, which only integrate into adult circuits during metamorphosis in the pupal stage^35^ (Fig. 1a). We identified primary and secondary populations in the atlas by the mutually exclusive expression of *Imp* and *dati*^36,37^ (Fig. 1d) and found they segregate into a region of punctate small groupings (“clusters”) and a region of elongated low-dimensional manifolds (“trajectories”), typically radiating from the “stem”, where neuronal precursors and undifferentiated neurons are located (Extended Data Fig. 3). Outside the small clusters, we found *Imp* expression at the tips of trajectories, distal to the stem. These are likely to be the first born neurons produced when the stem cell progenitors (neuroblasts) resume division after larval hatching and were annotated as “early secondaries”^38^. This neurogenesis wave annotation is supported by the observation that secondary neurons make up 59% of the transcriptional atlas, similar to the 60% reported in a recent scRNAseq larval atlas^38^ and to the 66% observed in the male VNC connectome (Fig. 1e). In addition, there is a similar trend in cell size, where primary > early secondary > secondary, as measured by number of RNA molecules per cell (transcriptome) and soma volume (connectome) (Fig. 1f).

The segregation of primary and secondary neurons into different regions of UMAP space highlights the neurogenesis wave as a strong driver of transcriptional divergence. However the local organisation is very different (punctate primary clusters versus elongated secondary trajectories) and suggests that these two populations may differ in the degree of separation between one cell type identity and its closest neighbour, with consecutively born primary neurons diverging more rapidly than secondary neurons. This dual organisation is preserved when the two subpopulations are analyzed independently, confirming that it is not a feature of the joint analysis, where higher diversity within primaries could mask diversity within secondaries (Fig. 1d). Strikingly, many primary clusters had sizes compatible with 1 cell, which is the median number of cells per type in the connectome (Fig. 1g), while the median cluster size is 3 cells. In contrast, secondary neurons yielded a median of 13 cells per cluster at the same clustering resolution, compared to 2 in the connectome (Fig. 1g). Moreover, secondary clusters exhibited greater overlap than primary clusters, with inter-cluster connections displaying an ordered, trajectory-like organisation in secondary neurons, while primary clusters had a less structured and more heterogeneous pattern (Fig. 1h). This observation aligns with connectomic analysis, where we find that primary neurons are statistically more distinct from their most similar neighbour than secondary ones (Fig. 1i) and that the nearest neighbour of a secondary neuron is more likely to have the same developmental origin (i.e. hemilineage identity) than for primary neurons (Fig. 1j). Together these results strongly support the idea that consecutively born secondary neurons diverge more gradually than primary neurons.

### Lineage identity links the atlas to the connectome

The existence of distinct progenitor pools in discrete spatial locations is another widespread feature of neural development. In the *Drosophila* VNC, secondary neurons are born from a defined set of larval neuroblasts (on average 25 per thoracic segment) each defining a stereotyped neuronal lineage. Sibling neurons from each lineage are further subdivided into hemilineages based on Notch signalling during asymmetric cell division: A (Notch ON) and B (Notch OFF)^33,39,4033,39^ (Fig. 2a). Crucially, neurons of the same hemilineage bundle together, leaving a structural footprint that has been used to group them in the connectome with >90% hemilineage annotation coverage^2^.

**Figure 2.**
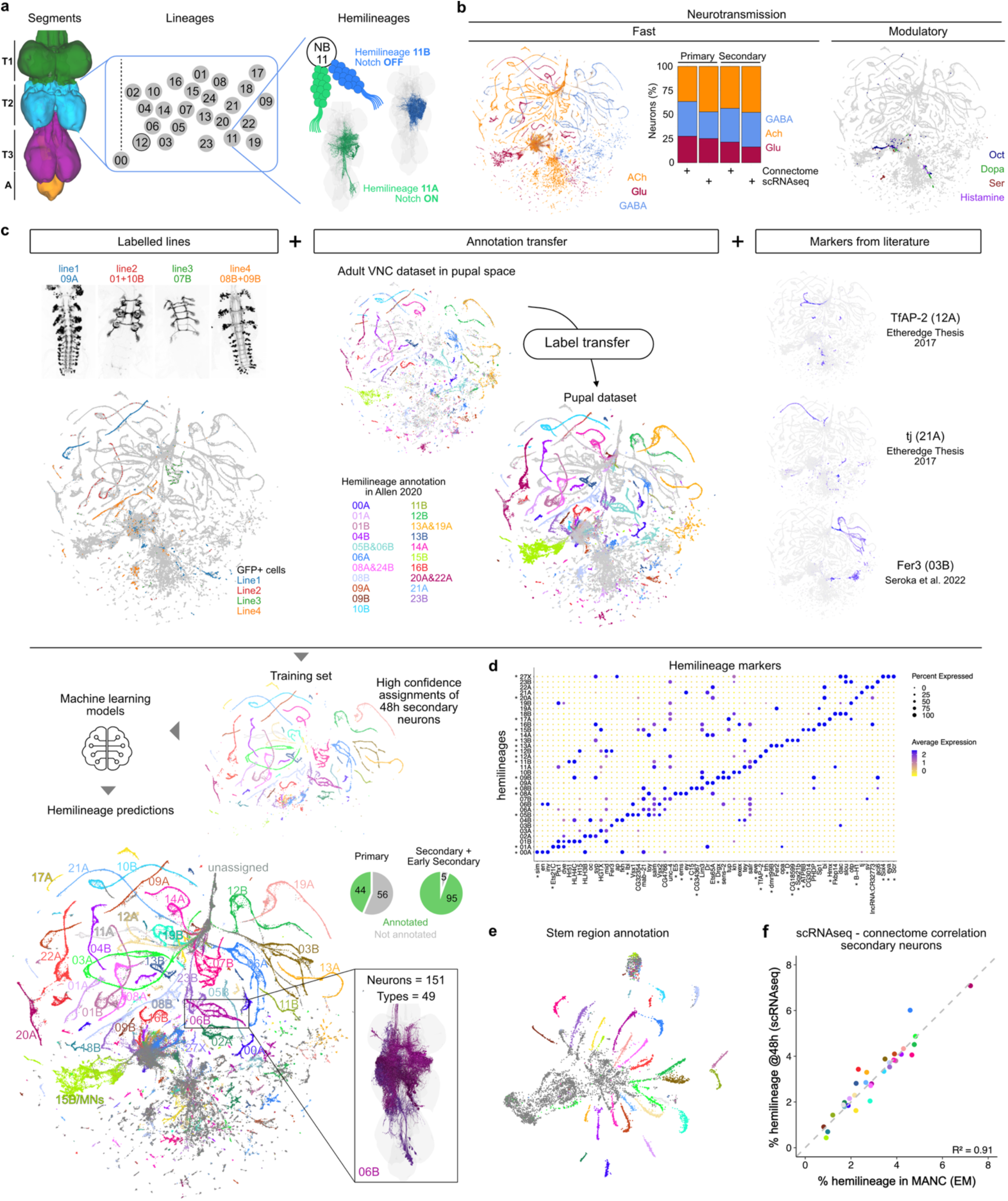
Hemilineage annotation links the transcriptome to the connectome. **a**, Adult VNC showing thoracic (T1, T2, T3) and abdominal A segments in green, blue, magenta and orange. Hox genes determining segmental identity are indicated. 25 Stem cells (neuroblasts, NB) in each thoracic segment generate a secondary lineage whose progeny is split in A and B hemilineages depending on Notch signalling. Renderings from the male connectome for hemilineages 11A and 11B. **b,** UMAP plots showing neurotransmitter predictions (fast and modulatory) based on expression levels of biosynthetic enzymes and transporters (see Extended Data Fig. 4). Fraction of fast neurotransmitter identities predicted in primaries and secondary+early secondary neurons in the atlas (48h) and the connectome. **c**, Pipeline to assign hemilineage identity. Top left: Maximum intensity projections of confocal stacks showing GFP expression pattern of Gal4 lines used for generating the atlas. GFP positive cells coloured by the genotype of origin. Top centre: adult VNC scRNAseq data from Allen et al.^31^ in the UMAP space of our atlas and the corresponding labels transferred to our atlas. Top right: markers from the literature used to de-orphanise remaining hemilineages. Middle: Assignments from Top used to train a machine learning classifier to predict hemilineage identity throughout the atlas. Bottom: Final hemilineage annotation. Rendering of MANC neurons from hemillineage 06B highlighting cross-matching between connectomic and transcriptomic atlases. Pie charts: fraction of annotated primary and early secondary+secondary neurons. MNs = motor neurons. **d**, Dot plot showing expression of hemilineage-selective marker genes in secondary neurons. Top 3 positive markers expressed with a log2FC > 0.5, in more than 70% of cells in the hemilineage and less than 20% of cells in the rest of the dataset. Asterisks indicate hemilineages with unique markers and the markers themselves. **e**, UMAP plot for precursor cells showing their manually curated annotation based on markers from (d). **f**, Correlation between % of secondary neurons per hemilineage annotated in EM and % of secondary neurons at 48h annotated in the atlas. Dotted grey line represents the identity line (y = x).

Previous work in adult VNC suggested that hemilineage identity is one of the main sources of transcriptional heterogeneity^31^. We therefore reasoned that the trajectories we observe in UMAP space could reflect the hemilineage organisation of the VNC. To address this, we first explored neurotransmission in our atlas, as neurons from the same hemilineage typically share neurotransmitter usage^2,41,42^. We found that cells in each trajectory expressed primarily just one fast-acting neurotransmitter (Fig. 2b, Extended Data Fig. 4a,b), supporting the idea that trajectories represent hemilineages. Aminergic modulatory neurotransmission is instead restricted to a small number of primary neurons (Fig. 2b, Extended Data Fig. 4c,d). Our atlas confirms the connectome predictions that across all neurons, excitatory acetylcholine is the most common fast neurotransmitter, followed by GABA and glutamate^2^.

To provide direct experimental evidence for hemilineage identity within our atlas, we included samples from four GFP reporter lines that label specific hemilineages (09A, 01+10B, 07B, 08B+09B)^43^ throughout development and into adulthood; distribution of GFP positive cells along specific trajectories confirms our prediction that trajectories correspond to hemilineages (Fig. 2c top left). To annotate hemilineages across the atlas, we combined multiple sources of direct and indirect evidence: labeled lines, annotation transfer from the adult VNC atlas^31^, a combination of markers from the literature, neurotransmitter usage and relative hemilineage sizes in the connectome (Fig. 2c top, Supplemental File 1 and Methods). We defined a set of high confidence assignments for 48h neurons and then trained a machine learning classifier to predict hemilineage labels across the atlas (Fig. 2c bottom). Predictions cover over 95% of secondary and 44% of primary neurons linking the hemilineages in our atlas to groups of, on average, 153 neurons/60 neuronal types in the connectome (e.g. 151 neurons/49 types in 06B). We then extracted markers for Notch status (Extended Data Fig. 5), which did not retrieve a global signature across lineages, and markers for hemilineage identity, 88% being transcription factors (Fig. 2d).

We found that 15 out of 34 hemilineages can be distinguished by the expression of a single marker, while the remaining ones can be identified using marker combinations; these markers should provide genetic access to hemilineages. Using these markers we extended hemilineage annotation to neurons in the stem region, which were poorly predicted by our automated classifier due to their immature state (Fig. 2e). The accuracy of our atlas annotation is supported by a very strong correlation between the observed sizes of hemilineages in the atlas and the connectome (R^2^=0.9, Fig. 2f).

### Segment annotation refines the link of the atlas to the connectome

Like the vertebrate spinal cord, the VNC has a repeated segmental organisation: homologous lineages are generated across segments, acquiring segment-specific characteristics through antero-posterior patterning specified by gradients of Hox gene expression^44,45^. We found evidence for this segmental organisation in the atlas: some hemilineages segregate into parallel trajectories with mutually exclusive Hox gene expression (Fig. 3a, Extended Data Fig. 6a,d). To refine our transcriptome-connectome matching, we defined Hox gene expression thresholds for each hemilineage-segment combination (Supplemental File 4). Hemilineage-specific thresholds were needed because Antp and Ubx antibody staining in L3 larvae revealed a more complex expression landscape than expected by the canonical view (Supplemental File 4). For example, Ubx protein levels were medium in T2 and high in T3 for 03B, but medium in T3 and high in A in 07B, reflecting expression level differences detected in the atlas (Fig. 3a,b, Extended Data Fig. 6b, Methods)^45^. In the 6hAPF dataset we also detected expression of the Hox genes *Scr* and *Dfd*, which are specific to the ventral region of the brain (the gnathal ganglia, GNG)^46^, likely resulting from imprecise cutting (Extended Data Fig. 6c,d). Accordingly, we annotated these neurons as GNG. In total, we provide segment identity predictions for 94% of secondary neurons and 43% of primary neurons (Fig. 3c).

The intersection of hemilineage and segment predictions subsets the atlas into 122 groups of secondary neurons, each mapping onto a mean of 51 neurons or 26 types in the connectome. The quality of the matching is supported by a strong correlation between the fraction assigned to each group in either dataset (R^2^=0.71, Fig. 3d).

For some hemilineages, intersegmental differences are large, resulting in independent trajectories each displaying specific marker genes (e.g. 03B), while others overlap partially (e.g. 01A) or completely (e.g. 13A) (Fig. 3e top and Extended Data Fig 7). We hypothesised that this difference in trajectory complexity could underlie the different degree of neuronal inter-segmental (serial) homology observed in the connectome (e.g. IN13B005 type has serially homologous neurons in all three thoracic segments Fig. 3e). To test this, we classified hemilineages into three categories of complexity: low (fully overlapping), medium (partially overlapping), and high (non-overlapping). We found that the transcriptional categories are significantly linked to the degree of inter-segmental homology reported in the connectome^2,47^ (Fig. 3e bottom). Interestingly, *Antp* is expressed across the three segments in low complexity hemilineages, instead of the expected restriction to T2 (Fig. 3f, Extended Data Fig. 6b), suggesting that Antp may contribute to reinforcing neuronal homology across thoracic segments. This might be needed to obtain neuronal identities implementing segmentally repeated sensory-motor circuits (e.g. leg movement).

**Figure 3.**
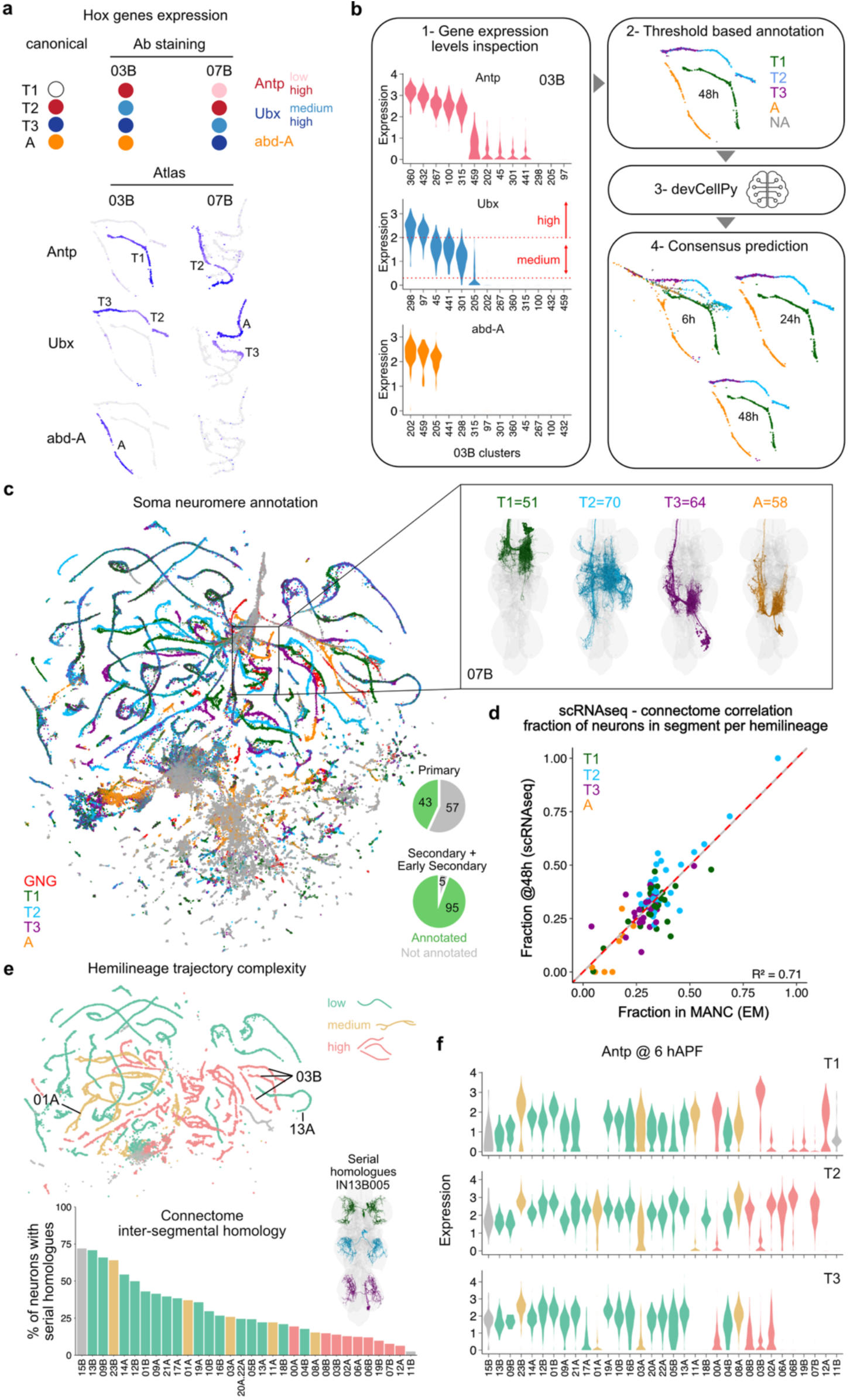
Segment annotation refines the transcriptome-to-connectome link. **a**, Top: canonical Hox genes (Antp, Ubx, abd-A) expression across segments and observed expression in 03B and 07B by antibody staining in larvae. Bottom: UMAP plots showing Hox gene expression in secondary neurons of 03B and 07B at 48h. The predicted segmental identity of the expressing trajectory is indicated. **b,** Pipeline to annotate soma segments. Left: violin plots showing per cluster expression in 03B secondary neurons at 48h. In 03B Ubx has a bimodal expression. Right: top, annotation based on Hox genes thresholds and used to train devCellPy classifiers (middle). The devCellPy model was used to predict soma segment identity across stages (bottom). **c,** Final soma segment annotation across all hemilineages and stages. Gnathal ganglia (GNG) segments (LB/MX/MD) are present only in the 6h dataset. Inset shows MANC 07B secondary/early secondary neurons (LHS only). Pie charts show the fraction of annotated primary and early secondary+secondary neurons. **d,** Soma segment annotation comparisons between atlas and connectome. Correlation between the fraction of secondary neurons per segment per hemilineage annotated in EM and that of secondary neurons at 48h annotated in the atlas. Dotted grey line represents the identity line (y = x). Solid red line shows the fitted linear model. **e,** Top: UMAP plot colour-coded by hemilineage trajectory complexity. Bottom: histogram showing, for each hemilineage, the proportion of secondary neurons with serial homologues in MANC. Hemilineages are ordered by decreasing inter-segmental homology. Trajectory complexity in the atlas correlates with inter-segmental homology in the connectome. Statistics: Jonckheere-Terpstra test, p-value = 2e-04. An example of a neuronal type with serial homologues across the three thoracic segments is shown. **f.** Violin plots showing Antp expression at 6h in the different hemilineages across segments and coloured by UMAP trajectory complexity. Hemilineage order as in (e).

To validate NM annotations we tested the expression pattern of the segment-specific markers *tey* (in 03B and 07B) and *kn* (in 07B) (Extended Data Fig. 7 and 8). As predicted, while in hemilineage 12A tey expression spanned T1, T2 and T3 segments, in 03B and 07B it was limited to the abdominal segment (Extended Data Fig. 8a-d). Expression of kn, instead, was limited to a small subset of T1 neurons and a large subset of T2 neurons in 07B, in agreement with our prediction (Extended Data Fig. 8e). Segment annotation is not perfect: while we predicted a very small number of tey+ 07B-T3 neurons, we did not detect any. As these cells arrange along an otherwise abdominal-specific trajectory, it is possible that they belong to the abdominal segment despite having lower Ubx levels.

### A global temporal code for neuron birth order

Sequential generation of different neuronal types from shared progenitors is a core developmental feature of the insect CNS^48–50^, but is also observed in other systems such as the mammalian retina^51^. Strikingly, we found strong evidence in our VNC atlas that neurons are arranged along trajectories according to their consecutive birth order from neuroblast progenitors: *Imp* is expressed at the tips of trajectories opposite to the stem (Fig. 1d) and previously reported genetic markers of birth order identity^52,53^ (*mamo*, *jim*, and *br*) are expressed in a conserved order along trajectories (Fig. 4a, Extended Data Fig. 9a). We reasoned that neuronal diversity within lineages should arise from changes in gene expression along each trajectory (Fig. 4a). To identify these genes we performed trajectory inference analysis: for each hemilineage-segment combination, we fitted a single curve and then calculated pseudotimes from first born neurons (pseudotime=0) to last born^54–58^ (Fig. 4b, Methods). We identified a total of 1,508 genes differentially expressed along the trajectories for all hemilineages, of which 149 were TFs (Fig. 4c). Strikingly, 40 of those TFs were common to more than 50% of the hemilineages, raising the possibility of a shared temporal program across hemilineages. This was supported by the observation that the relative order in which some TFs are expressed is conserved (Fig. 4d, Extended Data Fig. 9b). Importantly, although some of these TFs were also expressed in the stem region, their relative expression order was preserved in more committed neurons and invariant across developmental stages, even when assessed relative to bona fide cell-type-specific markers such as neuropeptides (Extended Data Fig. 9c).

**Figure 4.**
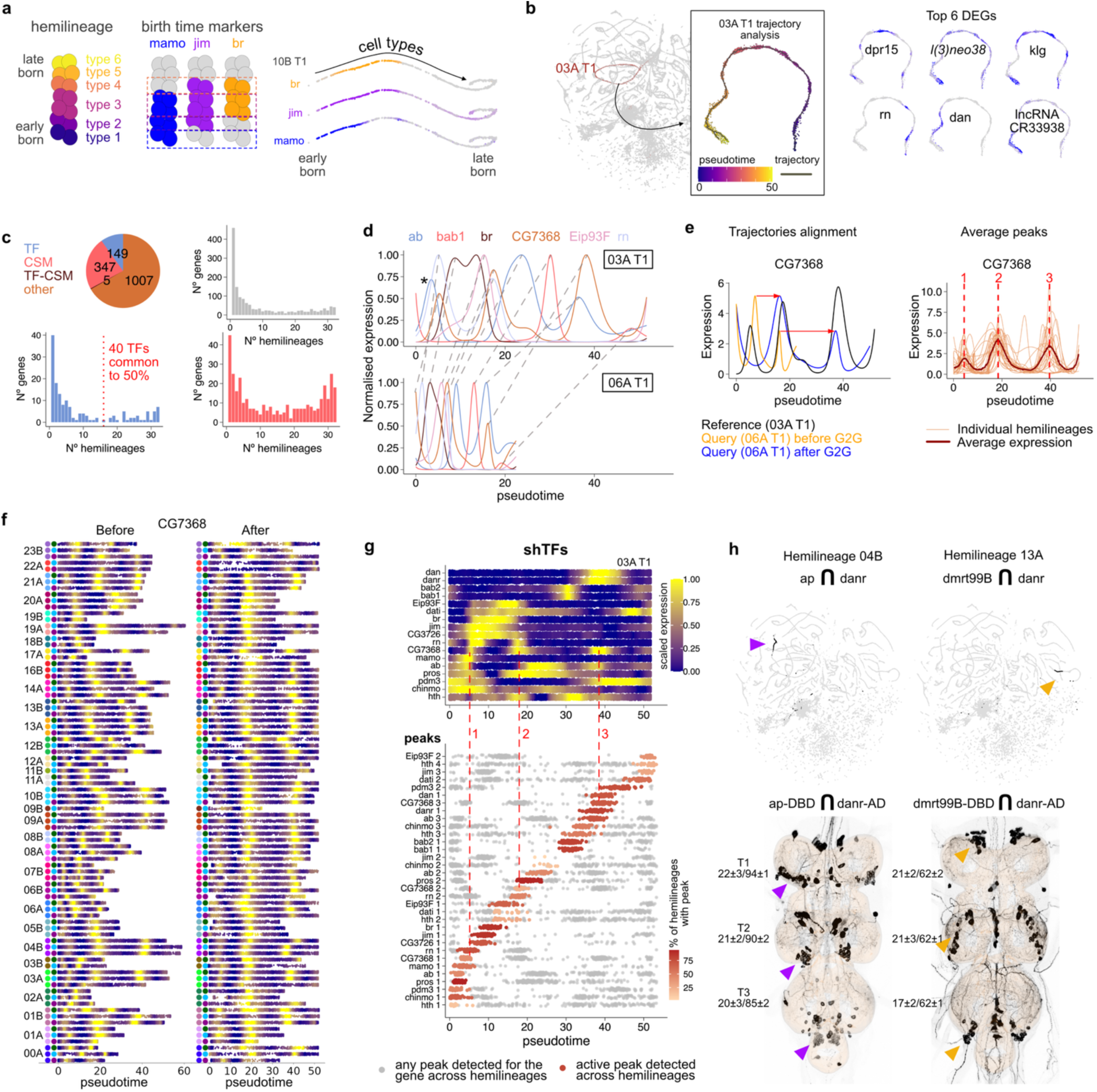
17 Transcription Factors expressed in a conserved temporal order across lineages define a persistent molecular correlate of birth order. **a**, Schematic illustrating cell types born sequentially within a hemilineage (left). UMAP embedding of hemilineage 10B-T1 forming a continuous trajectory where three genetic markers of birth order identity are expressed consecutively. **b,** Hemilineage trajectories analysis: pseudotime values from monocle3 for 03A-T1 trajectory and the 6 most significant differentially expressed genes (DEGs). **c,** Summary of DEGs along all trajectories. Pie chart of the distribution of gene categories across all analysed trajectories. DEGs with q-value < 0.01 and Moran’s value > 0.3 are shown. Histograms show the number of DEGs (y-axis) shared across hemilineages (x-axis), grouped by category. 40 TFs are shared by 50% of the 33 hemilineages analysed. Grey = all DEGs, blue = TFs, red = Cell Surface Molecules (CSM), brown = dual function. **d,** Fitted profile of normalised expression along pseudotime in 2 trajectories for a subset of TFs differentially expressed in 95% of hemilineages. Dashed lines link matching expression peaks. Asterisks indicate a peak present in 03A-T1, but absent in 06A-T1. **e,** Left: expression profiles of CG7368 along pseudotime for 06A-T1 before and after G2G alignment to the reference trajectory (03A-T1). Right: expression profiles of CG7368 along pseudotime after alignment in all trajectories. Average expression profile and average peaks are also shown. **f,** Heatmaps showing expression (normalised between q05 and q95) of CG7368 along pseudotime in all analysed trajectories before (left) and after (right) alignment. Colours in y axis: hemilineages (left, refer to Fig. 2) and segments (right, green = T1, blue = T2 and purple = T3). **g,** TFs expressed in a shared temporal order in all VNC lineages (shTFs). Top: heatmap showing normalised expression (between q05 and q95) along pseudotime of shTFs ordered by the first peak in the consensus sequence in 03A-T1. Bottom: gene expression peaks positions in pseudotime (aligned to reference) arranged by the consensus order. Each dot is a peak detected in a single trajectory. Peaks matching the average “active” peak are colored (intensity reflects detection frequency across trajectories); other peaks are shown in grey. **h,** Intersecting shTF *danr* with hemilineage specific markers labels specific neuronal subpopulations in adults. Top: UMAP showing cells co-expressing hemilineage markers (*ap* for 04B and *dmrt99B* for 13A) and *danr*. Bottom: adult VNCs showing split-GAL4 intersection patterns for the same genes. Max intensity projection of confocal stacks are shown. Images are registered to the VNCIS2 common template. Numbers: average neurons labelled per side belonging to the hemilineage ± SD / average number of neurons per side in the hemilineage in the connectome. Arrowheads point to neurons belonging to expected hemilineage. Additional smaller clusters not belonging to, respectively, 04B or 13A hemilineages are visible, likely primary neurons, in agreement with atlas prediction.

However, because pseudotimes were inferred independently for each trajectory, their scales are not directly comparable, even though they should all represent a monotonically increasing function of neuronal birth order. To facilitate a more direct comparison of gene expression dynamics across different hemilineage-segment trajectories and test whether more genes follow this common temporal order, we aligned all trajectories to a shared pseudotime space. This alignment assumes that, despite differences in local dynamics, a global temporal axis exists, reflecting a consensus sequence of transcriptional states. For this we used a recently developed nonlinear alignment method based on gene expression profiles^59^. We selected hemilineage 03A-T1 as the reference trajectory, as it is one of the longest, continuous trajectories in the atlas (Fig. 4e, Extended Data Fig. 9d,e). This approach enabled us to identify expression peaks that occur at the same ‘aligned’ pseudotime across hemilineages and segments (Fig. 4e,f). As a result, we found 17 TFs that exhibit a conserved pattern of expression along pseudotime and across hemilineages (*hth*, *chinmo*, *pdm3*, *pros*, *ab*, *mamo*, *CG7368*, *rn*, *CG3726*, *jim*, *br*, *dati*, *Eip93F*, *bab1*, *bab2*, *danr*, *dan*) (Fig. 4f,g, Extended Data Fig. 10). Given their shared expression dynamics across hemilineages, we refer to these as “shared TFs” (shTFs). Several shTFs are expressed at multiple points along the trajectories, expanding the total to 33 distinct expression peaks occurring in a consensus temporal order.

The combinatorial action of shTFs offers a possible mechanism to generate neuronal diversity. However, it also offers the promise of a modular genetic approach to access cell types, which could be of great experimental significance. We tested this possibility by using the split-GAL4 system to label cells in adult VNCs born during the *danr* temporal window from hemilineages 04B and 13A. We observed specific labeling of small neuronal populations within these lineages, recognisable by their morphologies (Fig. 4h). This result also indicates that *danr* expression is retained in adults. Like *danr*, most shTFs show persistent expression throughout developmental stages (Extended Data Fig. 9f). We confirmed protein expression in adults for 9 shTFs (Hth, Bab1/2, Dan, Br, Eip93F, Mamo, Pdm3 and Pros), in both VNC and central brain, revealing co-expression patterns compatible with transcriptional predictions (Extended Data Fig. 11).

To link shTFs expression patterns to neuronal birth-time we performed pulse-chase labelling, in which larvae were fed EdU during non-overlapping time windows (Fig. 5a). These experiments confirmed that neurons expressing two TFs mapped to non-overlapping temporal domains in the atlas (*br* and *bab1*, Fig. 5b,c) are born in distinct temporal windows in the larva: Bab1+ cells are born in early and late windows, while Br+ cells are born in between (Fig. 5d), consistent with the pattern in the atlas.

Unlike the well-characterized transcription factor cascades operating during embryogenesis in progenitor cells^4,60,61^, where temporal identity is transient and tied to progenitor state, the expression peaks defined by the 17 shTFs provide neurons with a persistent molecular timestamp, reflecting the time of neuronal birth.

### The shTF Br defines neuronal identity

The continuous expression of shTFs is compatible with a role as terminal selector genes^62^. We explored this possibility by testing the effect of ectopically expressing Br in hemilineage 14A. Since Br is normally present in a fraction of the hemilineage neurons, we asked whether extending its expression beyond its canonical temporal window would alter the fate of other 14A neurons. Using Bab1 expression as a readout of genetic interaction between shTFs expressed in non-overlapping temporal windows (Fig 5c), we found that while Bab1 is readily detected in control animals, ectopic Br eliminates strong Bab1 expression within 14A neurons, without affecting other lineages (Fig. 5e). Furthermore, there is a significant reduction of some projections and an enhancement of others (Fig. 5e), with no significant changes in the number of neurons (in T1 NM: OE = 152±1.5 SEM, n = 4; control = 155.3±2.0 SEM, n = 3; two-tailed t-test, *p* = 0.23). Interestingly, in the MANC connectome we identified 14A neuronal types compatible with the lost or gained projections (Fig. 5f). These results indicate that ectopic Br expression is sufficient to shift neuronal identity within the 14A hemilineage, supporting its role as a terminal selector in the VNC.

### The temporal transcription factor code is conserved in the brain

To explore if the shared transcription factor code extends to secondary neurons in the brain we analysed a small subset of hemilineages in two brain regions: the GNG, evolutionarily closely linked to the VNC, and the Superior Lateral Protocerebrum, a more derived region without evident transcriptional, morphological or functional homology with the VNC^63^ (Fig. 5g). For the GNG, we took advantage of trajectories present in our 6h samples due to imprecise separation of brain and VNC tissues (Extended Data Fig. 6c,d), while the Superior Lateral Protocerebrum hemilineage was isolated from an independent brain dataset (see Methods for data source).

**Figure 5.**
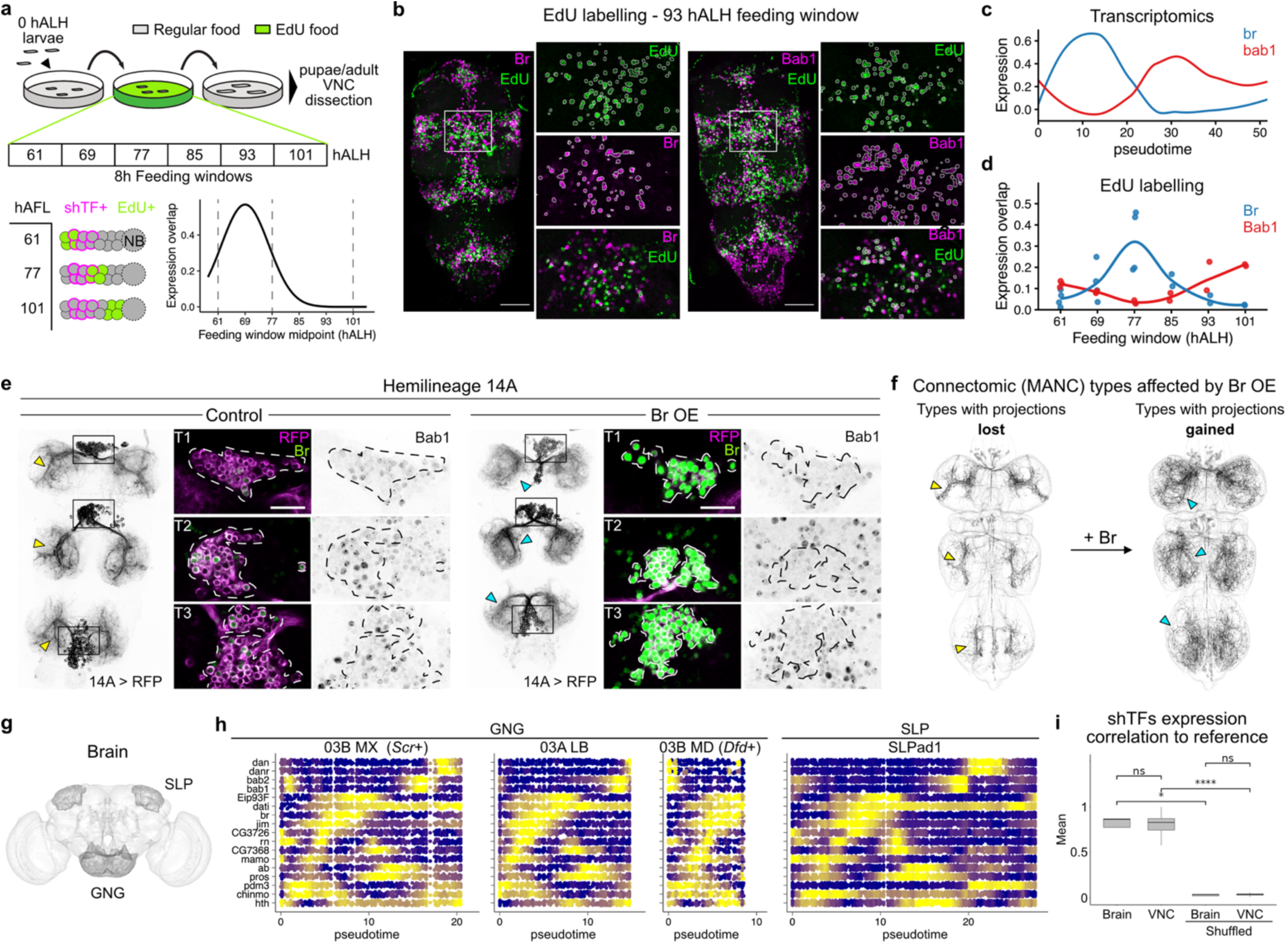
The VNC’s global temporal transcription code is also conserved in the brain. **a**, Schematic of EdU pulse-chase experiments. **b,** Images show representative examples of EdU-labelled cells (green) in larvae fed during the 93 hALH feeding window, together with the shTFs Br (magenta, left) or Bab1 (magenta, right). Whole VNCs are displayed as MIPs. Insets show a single z-plane and the corresponding segmentation results for EdU (top), shTF (middle), and their intersection (bottom). Scale bar: 50 µm. **c,** Expression levels along pseudotime for br (blue) and bab1 (red) in 03A-T1 lineage. **d,** Fraction of shTF+ cells colocalising with EdU across feeding windows (see Methods, EdU labelling). Each dot represents one VNC (n=2 to 4/condition). **e,** Overexpression (OE) of Br-Z4 isoform in 14A hemilineage causes the disappearance of Bab1+ cells within 14A and a change in projections morphology. Projections lost (yellow) or gained (cyan) in Br-Z4 OE condition are marked by arrowheads. Whole VNCs are displayed as MIPs. Insets show a single z-plane. Scale bar: 20µm. Images are representative of n=4/condition. **g,** Schematic of the brain highlighting regions of origin for the hemilineages examined in (h). **h,** Heatmaps showing the expression of shTFs for hemilineages in the brain as in Fig. 4g top. GNG data from 6h and superior lateral protocerebrum (SLP) data from 6-48h. LB = labial, MX = maxillary, MD = mandibular. **i,** Mean correlation of shTFs expression between the reference trajectory (03A-T1) and aligned brain trajectories from (h) or VNC ones. Box-and-whisker plot shows the median and interquartile range across trajectories. For the shuffled condition shTF identity labels were shuffled and correlations calculated 200 times. Statistical test: Wilcoxon rank-sum test.

In both regions and across all four hemilineages, shTFs exhibited expression patterns consistent with those observed in the VNC (Fig. 5h). We quantified this observation by calculating expression correlation between each hemilineage and the 03A T1 reference. We did not observe a statistically significant difference between brain and VNC scores (Fig. 5i), supporting the notion that shTFs influence the development of all secondary neurons in the central nervous system, excluding the optic lobes. These results describe a universal transcriptional code that links birth order to neuronal identity.

### Mapping sex differences across the atlas and the connectome

Many aspects of sexually dimorphic behaviours are encoded by neural circuits in the VNC. For example, males vibrate their wings to produce a courtship song^64,65^, while females become less receptive and lay more eggs after mating^66^.

The existence of both male and female VNC connectomes enables comparative analysis between sexes and the identification of dimorphisms at scale^20^. Integrating connectomic datasets with molecular profiles across development offers a powerful framework to investigate the origin and nature of sex-specific neuronal differences.

While acquiring data, we were careful to sample a balanced number of female and male cells, whose sex could be unambiguously inferred from the genotype of the source animal (see Methods). Thanks to this, we were able to investigate which regions in the atlas displayed numerical biases in male or female neurons^67^. At 48h - the most mature stage in our dataset - we found that 4% of neurons belong to numerically biased regions, i.e. differentially abundant (DA) (Fig. 6a,b, Extended Data Fig. 12a). Male-enriched cells are more likely to be secondary neurons (81% of male-enriched neurons are secondaries), while female-enriched cells are biassed towards primary neurons (78% of female-enriched are primaries) (Fig. 6b). These differences were accompanied by selective enrichment of the key sex-determination genes, *fruitless* (*fru*) and/or *doublesex* (*dsx*)^68^, again biased by neurogenesis wave (Fig. 6b).

To link transcriptomic predictions to the connectome we focused our analyses on secondary neurons, given their more complete annotation by hemilineage and segment. Specific hemilineage–segment combinations emerge as hotspots of sexual dimorphism (Fig. 6c), including previously characterised dimorphic lineages, such as courtship song neurons in 12A-T1 and 08B-T1^20,64,65^. To validate our predictions using the male (MANC) and female (FANC) connectomes, we focused on 6 hemilineage-segment combinations: 2 with pronounced male enrichment (08B-T1 and -T2), one with known sexually dimorphic neurons^69^ (01A-T1), and 3 without significant sex-bias (01A-T2, 06A-T1 and 06B-T1) (Fig. 6c). Since no comprehensive one-to-one matching between MANC and FANC connectomes existed, we first established an automatic pipeline to align these datasets (Fig. 6d). We then transferred hemilineage labels from the well-annotated MANC to FANC, which has limited annotation and is only partially proofread. After proofreading and manually curated annotation of the selected FANC hemilineages (Methods) we found, as predicted, the greatest numerical differences between sexes in 08B-T1 and -T2, which also had the largest number of MANC-specific neuronal types (Fig. 6e).

**Figure 6.**
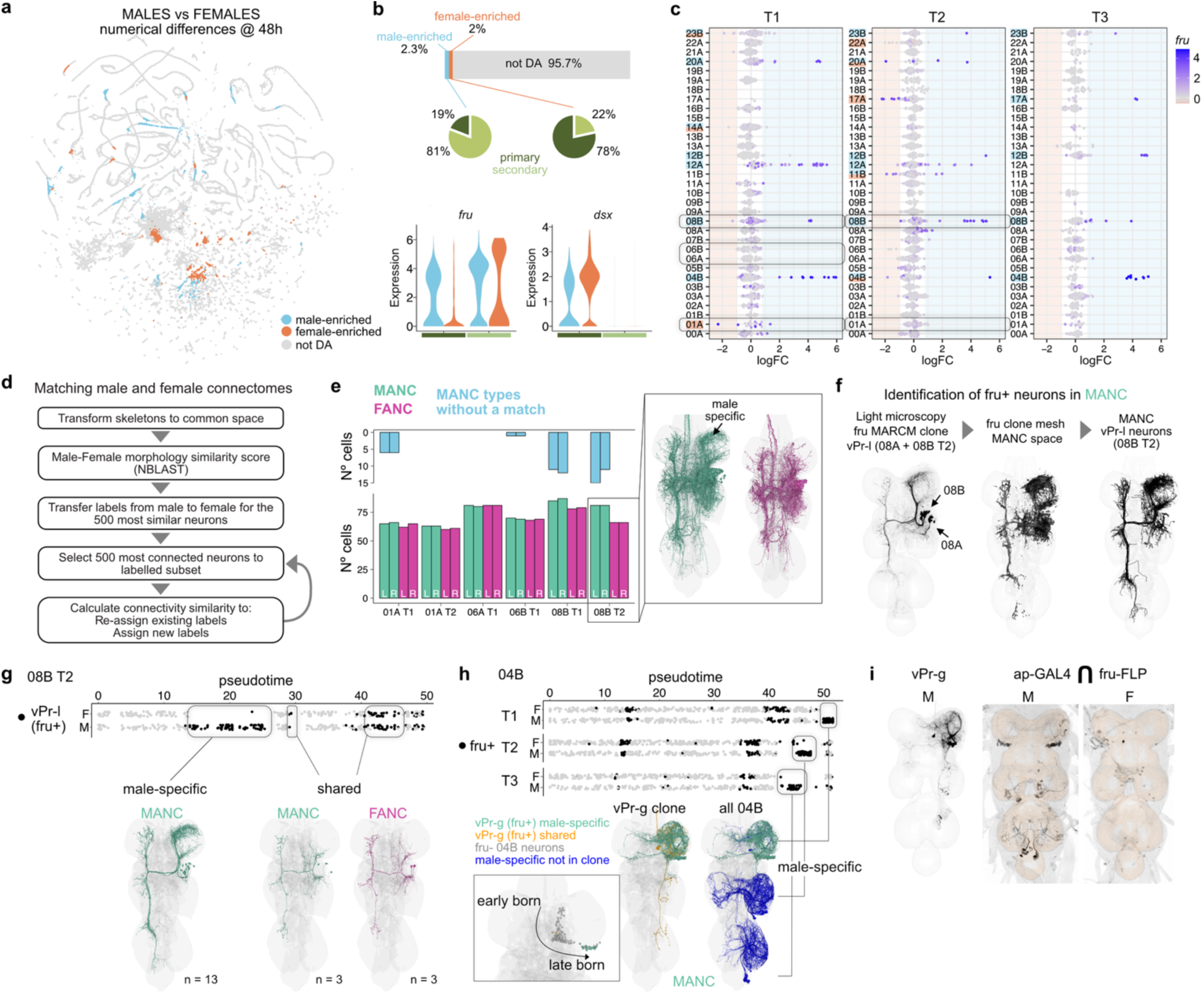
The transcriptional atlas predicts sex differences between male and female connectomes. **a**, Differential abundance (DA) between males and females at 48h. Cells in the UMAP plot are coloured by differential neighborhood enrichment (light blue = male-enriched, orange = female-enriched). **b,** Sex differences in primary and secondary neurons. Bar shows the % of neurons enriched in males, in females and not enriched at 48h. Pie-charts show the % of primary and secondary neurons in male-enriched or female-enriched neurons. Violin plots show expression of fru and dsx in sex-specific neighborhoods at 48h. **c,** Beeswarm plot showing differences in secondary neuron abundance at 48h, grouped by hemilineage and soma segment. Each dot corresponds to a neighborhood in milo analysis, coloured by the mean expression of fru. Boxes highlight hemilineages object of manual curation in FANC. **d,** Pipeline for matching neurons in EM between sexes. **e,** Hemilineage annotation in FANC. Lower histogram summarises the number of neurons identified per hemilineage, per side, per sex after manual curation (L= left, R = right). Upper histogram shows the number of MANC neurons belonging to types for which zero matches in FANC were found by the automatic matching pipeline. Inset shows neurons belonging to 08B hemilineage in T2 in MANC (green, L) and FANC (magenta, R) after manual curation of FANC in MANC space. **f,** Pipeline to identify candidate fru+ neurons in MANC starting from MARCM clones. **g,** The atlas predicts sex-specific and shared fru+ neurons in the connectome. Top: dot plot showing atlas secondary 08B-T2 neurons at 48h arranged by pseudotime and coloured by vPr-l fru clone. Bottom: neurons corresponding to vPr-l clone neurons in the connectome. Colours indicate the dataset of origin (MANC = green, FANC = magenta). Male-specific neurons are plotted on the left, shared neurons plotted on the right. Numbers indicate the number of plotted neurons. **h,** The atlas predicts fru+ male-specfic neurons in hemilineage 04B across segments. Top: dot plot showing atlas secondary 04B neurons at 48h in all segments arranged by pseudotime and coloured by fru expression. Bottom: vPr-g clone neurons in MANC (left). Colours indicate male specificity (green = specific, orange = shared). All male-specific 04B neurons in MANC (right). vPr-l neurons are shown in green, all other male-specific neurons in blue (neuron type not found in FANC by automatic matching). Inset shows soma position of 04B secondary neurons in T1 arranged according to birth order. **i,** Neurons labelled by genetic intersection of ap-GAL4 and fru-FLP in male (middle) and female (right), corresponding to 04B fru+ neurons. Max intensity projections of confocal stacks are shown. Male fru clone mapping to 04B-T1 is shown as reference (left).

In line with previous findings, nearly all sex-biased secondary neurons expressed *fru* (Fig. 6c), although not all fru+ neurons belonged to numerically dimorphic groups (Extended Data Fig. 12d). To map fru*+* dimorphic neurons from the atlas to the connectome, we identified them in MANC using a previously published dataset of light-level *fru* lineage-related neurons (MARCM clones)^69^ (Fig. 6f, Extended Data Fig. 13a,b, Methods, Supplemental File 2). Since this resource links gene expression, hemilineage identity and neuronal morphology, we were able to match precisely defined small groups of fru+ neurons across the atlas and the connectome.

We focused on fru+ neurons from the dimorphic hemilineage 08B-T2, which we mapped to the vPr-l fru+ clone. Pseudotime analysis shows that fru+ vPr-l neurons in the atlas are born in multiple time windows: an earlier born, larger cluster contains male-specific neurons, while 2 later born, smaller clusters have neurons from both sexes (Fig. 6g). In line with this prediction, the connectome revealed that most vPr-l types had no female counterparts (male-specific); only two types (three neurons per side) were shared (Fig. 6g). As a second example, in hemilineage 04B the atlas predicted male-specific neurons in all three thoracic segments (Fig. 6h), although a corresponding clone had previously been reported only in T1 (vPr-g)^69^. We confirmed the atlas’s prediction by using our MANC-FANC automatic matching and intersectional labelling experiments (Fig. 6h,i), which identified male-specific fru+ neurons in all three segments. Remarkably, our atlas clearly identifies the fru+ neurons as very late born, matching the consistent external position of the cell bodies of male-specific vPr-g 04B neurons in the MANC connectome (Fig. 6h).

These results showcase the atlas’s power for studying known and previously unknown sexual dimorphisms.

### Female-specific apoptosis and transcriptional divergence shape sexual dimorphisms

We next asked how sex differences in the VNC emerge during development. At 6h we observed minimal numerical differences across the atlas (compare Extended Data Fig. 12e and Fig. 6c), indicating that dimorphisms develop progressively throughout metamorphosis. Accordingly, in early stages there are neurons in both sexes within the vPr-l male-specific temporal window, but these are gradually lost through pupal development. This correlates with increased expression of the pro-apoptotic gene *rpr* at 24h only in females (Fig. 7a, see Extended Data Fig. 12f for a second example), suggesting that female-specific apoptosis is the reason why vPr-l neurons are male specific. The same observation was replicated across the atlas (Extended Data Fig. 12g), pointing to a widespread role for female-specific apoptosis in shaping male-specific neuronal domains, extending earlier results in the brain^70–72^ to molecularly defined cell populations.

While some neurons are sex specific, others only show morphological differences^69^. To investigate the molecular signatures underlying morphological dimorphisms we focused on hemilineage 01A-T1, where we mapped previously described sex differences in the dPr-a fru+ clone^69^ to both MANC and FANC connectomes (Fig. 7c). These dimorphisms are so pronounced that MANC neurons fail to match with their corresponding dimorphic FANC counterparts in Fig. 6a, despite a similar number of neurons per hemilineage in both sexes. The atlas shows that fru+ neurons display a clear separation between males and females in 01A-T1 UMAP space (Fig. 7d). We calculated gene expression differences between sexes in clusters of the 01A-T1 lineage and identified the dimorphic cluster to have the highest number (Fig. 7d). Interestingly, among the most significant genes we found several with synaptic partner matching functions, such as members of the *beat*, *side* and *dpr* families^73^ (Fig. 7e), that might play an instructive role in dimorphic wiring. Finally, we quantified the emergence of dimorphism through metamorphosis, by defining a per-cell male-female molecular distance metric (Fig. 7d, Methods). This shows that molecular distance, and the number of differentially expressed genes, increase with developmental time specifically in fru+ neurons (Fig. 7f).

Overall these results suggest that sexual dimorphisms emerge through development from neurons with the same temporal origin by employing two alternative strategies: sex specific apoptosis and transcriptional divergence. Two recent independent studies reached similar conclusions that sexual dimorphisms reflect sex-specific developmental programs acting on neurons born in the same lineage and birth order, rather than de novo generation of sex-specific cell types^74,75^.

**Figure 7.**
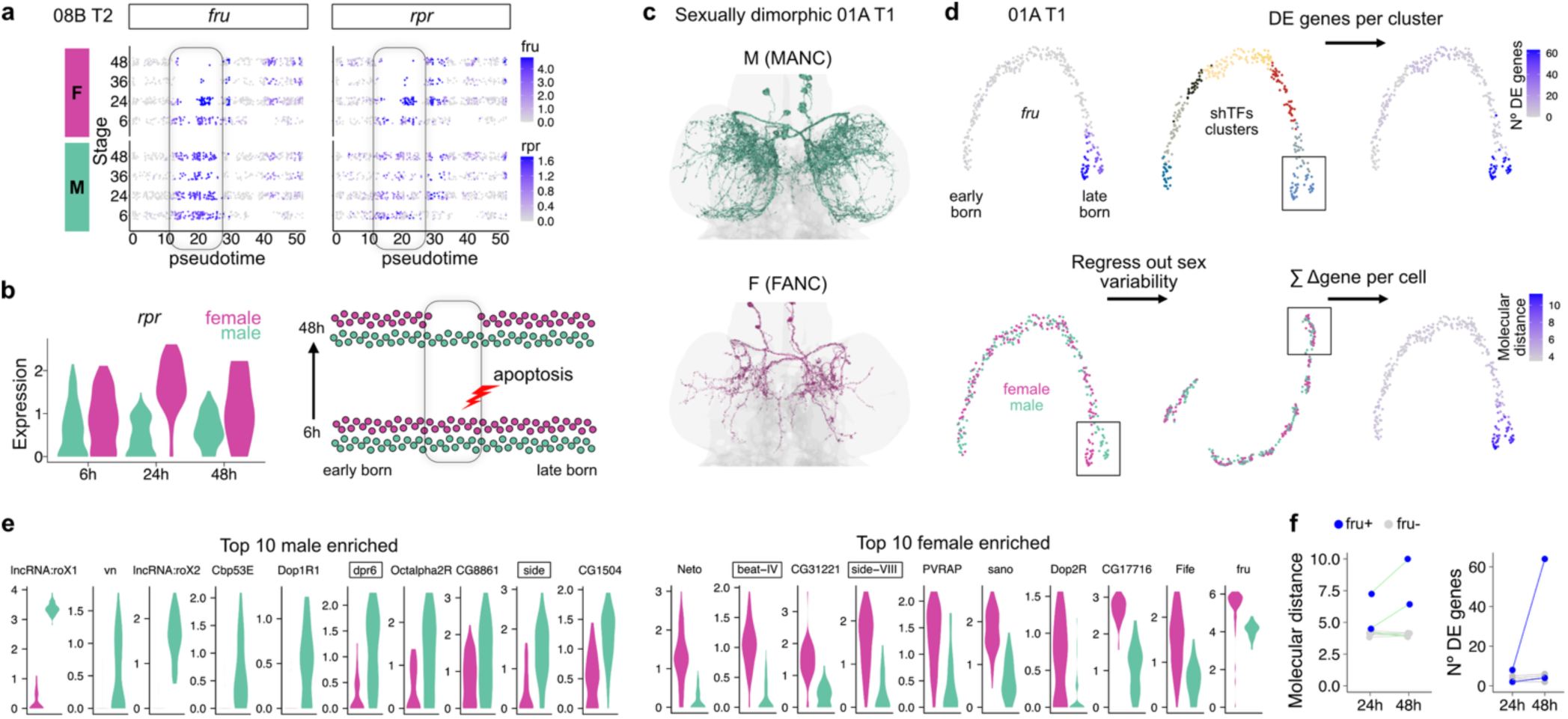
Male-specific apoptosis and transcriptional divergence shape sexual dimorphisms through development. **a**, Dot-plot showing expression of *fru* (left) and *rpr* (right) in 08B-T2 secondary neurons arranged along pseudotime and split according to sex and animal stage. **b,** Expression of *rpr* in the pseudotime window corresponding to male enriched neurons. Schematic of female specific apoptosis resulting in male-specific survival of birth-order-defined neuron types. **c,** Male and female sexually dimorphic fru+ neurons. **d,** UMAP plots showing 01A-T1 secondary neurons in the atlas at 48h. Cells are coloured by *fru* expression, cluster identity calculated using shTFs, number of DEGs per cluster, sex and per-cell correction score after regression of sex-identity. **e,** Expression of male and female enriched genes in the most dimorphic cluster at 48h. **f,** Molecular differences at 24 and 48h. Left: average correction score per cluster. Lines connect corresponding clusters, green for significantly different scores (t-test with Bonferroni correction). Right: number of DEGs between male and female per cluster at 24 and 48h. Blue dots = fru+, grey dots = fru-.

## Discussion

We report the first high resolution developmental transcriptional atlas of the *Drosophila* nerve cord. This provides a platform to investigate the molecular logic of the thousands of cell types that wire up to form the sensory-motor circuits in the nerve cord, similarly to how more focussed atlases within the brain have been used to understand wiring specificity of *Drosophila* olfactory^76^ and visual circuits^77–79^. We linked the atlas and the connectome by sequentially annotating primary and secondary neurons, hemilineages, segments and *fru* identity. Our comparative analysis of sexual dimorphisms in fru+ neurons across datasets led to a further refinement of cross-matching down to as few as 3 neurons (Fig. 6g). While our atlas has not yet been exhaustively annotated at the connectomic neuronal type level, the correspondence obtained for fru+ neurons shows that the dataset contains information at the necessary granularity to identify them. By describing the molecular identities of consecutively born secondary neurons as ordered trajectories, we open the possibility to align transcriptomic and connectomic types. Ranking neurons in the connectome by their birth order would naturally position them along these trajectories, facilitating the assignment of connectomic cell types to transcriptional profiles. This ranking could be established, as done here for *fru*, through stochastic labelling of neuronal morphology^48,49,80^, or through dense labelling of shTFs coupled with expansion microscopy-based neuronal reconstruction (LICONN)^81^.

Our data show that neuronal types born consecutively from the same progenitor in the larva diverge more gradually than those born in the embryo. This is supported by a striking difference in organisation within transcriptomic (UMAP) space: punctated small clusters vs elongated low-dimensional manifold trajectories. Interestingly, for some hemilineages, secondary neurons segregate into multiple trajectories, likely corresponding to distant anatomical subgroups. While trajectories do not break apart into well separated clusters, local thickenings exist which could house the most abundant cell types (see Extended Data Fig. 7b). Increasing the coverage of the atlas will establish whether each connectomic secondary type can resolve in an isolated cluster. While the distinction between primary and secondary neurons could reflect differences in maturation, with primary neurons being born earlier, three lines of evidence argue against it: firstly, the time difference between the birth of primaries and that of early-secondary neurons is smaller than between the latter and late-secondary neurons; secondly, similar differences between Imp+ and Imp-cells are also present in a recently published atlas of the adult CB (ref. to Fig. 2d and Fig. 2k in Allen et al. 2025); and, thirdly, we also detect corresponding morphological and connectivity differences in the adult connectome. These observations imply that UMAP structure reflects fundamentally different developmental programs rather than degrees of maturation. Studies in *Drosophila* may therefore help elucidate the developmental logic of contiguous neuronal types, previously associated with mammalian nervous systems^8,18,82^.

Notably, the temporal patterning mechanisms specifying embryonic-versus larval-born neurons mirror this distinct organisation: a discrete temporal cascade of TFs in embryonic neuroblasts^4,60,61^ versus opposing gradients of the RNA binding proteins Imp (early) and Syp (late) in larval neuroblasts^37,83^. The relationship between temporal origin and molecular identity has been characterized in embryonic and optic lobe neurons^4,84^, but how Imp/Syp gradients translate to neuronal identities is only starting to be explored^37,52,85^. We propose that the 17 shTFs are perfectly poised to be key intermediaries in this process. While the precise functional contributions of most shTFs in specifying and maintaining neuronal identity remain unresolved, instructive roles in fate specification have been uncovered for some of them^26,74,86,87^. Our Br-Z4 overexpression results provide an additional example of this role in the VNC. From an evolutionary perspective, a gradual molecular identity system could offer several advantages. It is more evolvable, as small shifts in gene regulation or timing can generate new neuron types without requiring entirely new patterning modules, enabling gradual circuit diversification and behavioral adaptation. It may also confer greater developmental robustness and plasticity, allowing neurons to adopt flexible wiring strategies based on graded molecular cues rather than fixed identities. In insects, which have a strongly constrained ground plan based on neuroblast lineages highly conserved across hundreds of millions of years of evolution^18,88^, this flexibility may be crucial to allow diversification of neurons and circuits across sexes and species.

The shTFs identified here are reminiscent of “concentric genes” that demarcate successive cohorts of neurons in the *Drosophila* optic lobe^22,84,89,90^. Despite this similarity, the shTF code appears more global as it is expressed across all the VNC hemilineages responsible for making thousands of cell types, regardless of Notch expression. The global scope of our atlas provides the strongest evidence to date for a persistent molecular correlate of birth order operating across secondary neurons of both the central brain and ventral nerve cord, extending temporal logic beyond previously studied, region-specific systems^22,32,52,84,89,90^. Consistent with our findings, nine shTFs had previously been identified as markers of specific projection neuron (PN) types, a lineage in which birth-order–dependent type specification has been extensively characterized^32^. In this context, *br* and *Eip93F* label mid-born PNs, whereas *danr* marks late-born PNs. In addition, two recent studies have identified most shTFs as temporally patterned in a small subset of central brain hemilineages in pupae and adults^74,91^. We find that additional transcription factors identified in these studies are not broadly shared across lineages in the VNC and therefore do not meet our criteria for inclusion among shTFs (Table 3).

Importantly, the temporal specification of neurons in flies has parallels in mammals. On the one hand, homologues of Imp and Syp exist and have been implicated in neurogenesis^92,93^. On the other hand, a similar shared transcriptional code has recently been described in maturing neurons of the mouse spinal cord, where early neurons express Onecut family TFs, intermediate neurons express Pou2f2, Zfhx3, and Zfhx4, and late-born neurons express Nfia and Nfib, often in combination with Neurod2 and Neurod6^94,95^. This evidence points to an evolutionarily conserved temporal logic governing the generation and organisation of neuronal diversity across species.

Given that this shared code is likely to result from the common evolutionary origin of progenitors, its conservation suggests functional importance. For example, shared temporal modularity could ensure stepwise circuit assembly and wiring coordination in local neuronal domains, as shown for local premotor circuits^96^, or between distant brain and VNC regions. This offers a mechanistic explanation and opens avenues of study for the idea that “neurons that are born together wire together”^97^. It may also confer shared functional properties, such as axonal targeting or circuit integration, to neurons born within specific temporal windows. The recurrent expression of the same TF in neurons from different temporal windows could be responsible for the previously noticed re-occurrence of morphological features in temporally distant neurons within lineages^48,49^. Interestingly, six of the shTFs we identified have established roles in regulating the specification of distal structures during appendage development (hth^98^, dan, danr^99^, bab1, bab2, rn^100^). Proximal-distal patterning has a temporal component, with positional identity being the result of the timing of exposure to spatial patterning factors. This suggests that the same regulatory networks may be repurposed in different developmental contexts or that there exists a deep common evolutionary origin for neuronal and non-neuronal tissue patterning.

We believe that many additional insights remain latent in the atlas, which will be a resource of value not only to the *Drosophila* neuroscience community, but also for comparative studies exploring the evolutionary origins of neuronal diversity and function across phyla. It also provides a strong case to obtain a correspondingly high coverage developmental brain atlas to complete the molecular description of the fly CNS.

## Acknowledgments

We thank the Cancer Research UK (CRUK) Cambridge Genomics Facility, particularly K. Kania, for generating and sequencing 10x libraries. We are grateful to members of S. Aerts lab: J. Janssens for initial advice on genetic demultiplexing and K. Davie for support with SCope. We acknowledge the contributions of Bucknell University undergraduates (K. Bell, G. Freehling, C. Murphy, A. Panzarino, A. Patterson and E. Schuler) for generating and analysing light-level Hox gene data, and J. Truman, in whose laboratory these experiments were initiated. We thank J. Phelps for assistance with the FANC dataset. We thank A. Schildkamp (Aelysia Ltd.) for assistance with hemilineage seeding and proofreading. We also thank L. Luo, G. Rubin, J. Parrish and S. Goodwin for sharing fly stocks and C. Desplan, N. Konstantinides, D. McKay, Muriel Boube-Trey for sharing antibodies. We are grateful to members of the Jefferis group, as well as R. Benton and N. Konstantinides, for comments on the manuscript.

This work was supported by an ERC Consolidator Grant (649111), core funding from the UKRI Medical Research Council (MC-U105188491), NSF NeuroNex2 (2014862) and AstraZeneca BlueSky (BSF2-16) to GSXEJ, as well as a Sir Henry Wellcome Postdoctoral Fellowship (110232/Z/15/Z) to E.D..

## Authors contributions

Conceptualization, S.C., E.D., and G.S.X.E.J.; methodology, S.C., E.D., and G.S.X.E.J.; writing, S.C., E.D., and G.S.X.E.J.; investigation, S.C., E.D., M.M., L.M., I.R.B., M.G., J.H.M.S.; resources, S.C., E.D., I.R.B., L.S. and H.L.; supervision, G.S.X.E.J., E.D. and S.C.

## Methods

### scRNAseq data generation

To discriminate technical from biological variability we used flies from different genetic backgrounds ^24,101^, enabling us to generate scRNAseq libraries of VNCs of different sex and developmental stages and separate them after sequencing through SNP-based sample demultiplexing (see below) ^102^ (Extended Data Fig. 1).

### Dissociation

For each stage-sex condition several genotypes and individuals were used, as summarised in Table 1. Flies of the genotypes indicated in Table 2 were allowed to mate and their larvae developed at 25°C until white pupae stage. Then pupae were collected with forceps, sexed under a dissecting scope and put in either male or female vials to develop until the correct stage, i.e. 6, 24, 36 or 48 hours after puparium formation (h). Ventral nerve cords (VNCs) were dissected in ice-cold DPBS (Dulbecco’s Phosphate Buffer Saline, Gibco). A maximum of 8 VNCs, each of a different genotype, were pooled in a low-bind eppendorf tube containing 100µl of DPBS on ice. VNCs from animals of two or three different developmental stages and sex were pulled in the same tube (6+24h, 24+48h or 24+36+48h stages). Four brain samples were included in the last round of experiments, but later excluded from the analysis of VNC. After dissections, tissues were collected at the bottom of the tube by centrifugation (800g, 5 mins, 4°C), DPBS was replaced with 50µl of dispase I (3mg/mL, Sigma-Aldrich, Cat. D4818, reconstituted in 50mM HEPES/KOH pH 7.4, 150mM NaCl) and 75µl of collagenase I (100mg/mL, Invitrogen, Cat. 17100-017, reconstituted in HBSS). VNCs were dissociated in a thermo mixer (Eppendorf) at 25°C at 500rpm for 30 (samples with 6+24h stages) or 40 mins (samples with 24+36+48h stages). Dissociation was aided by pipetting up and down 10 times every 10 mins ensuring that the tip touched the bottom of the tube to increase the mechanical stress. Dissociated cells were centrifuged (800g, 5 mins, 4°C), washed once with DPBS and resuspended in 180µl of DPBS with 0.04% BSA. Cell suspension was filtered through a 10µm filter (pluriStrainer, Cambridge Bioscience, Cat. 43-50010-03). Viability, presence of debris and cell number were accessed by a Countess II automated cell counter (Invitrogen).

**Table 1.** Summary of samples used to build the atlas. Stage in hours after puparium formation. Genotypes, number of different genotypes used per stage-sex condition. Independent samples correspond to the number of individuals used per condition.

| Stage | Sex | Genotypes | Independent Samples |
| --- | --- | --- | --- |
| 6 | F | 7 | 8 |
| 6 | M | 9 | 10 |
| 24 | F | 11 | 18 |
| 24 | M | 11 | 16 |
| 36 | F | 7 | 7 |
| 36 | M | 5 | 5 |
| 48 | F | 11 | 16 |
| 48 | M | 11 | 14 |

**Table 2.**
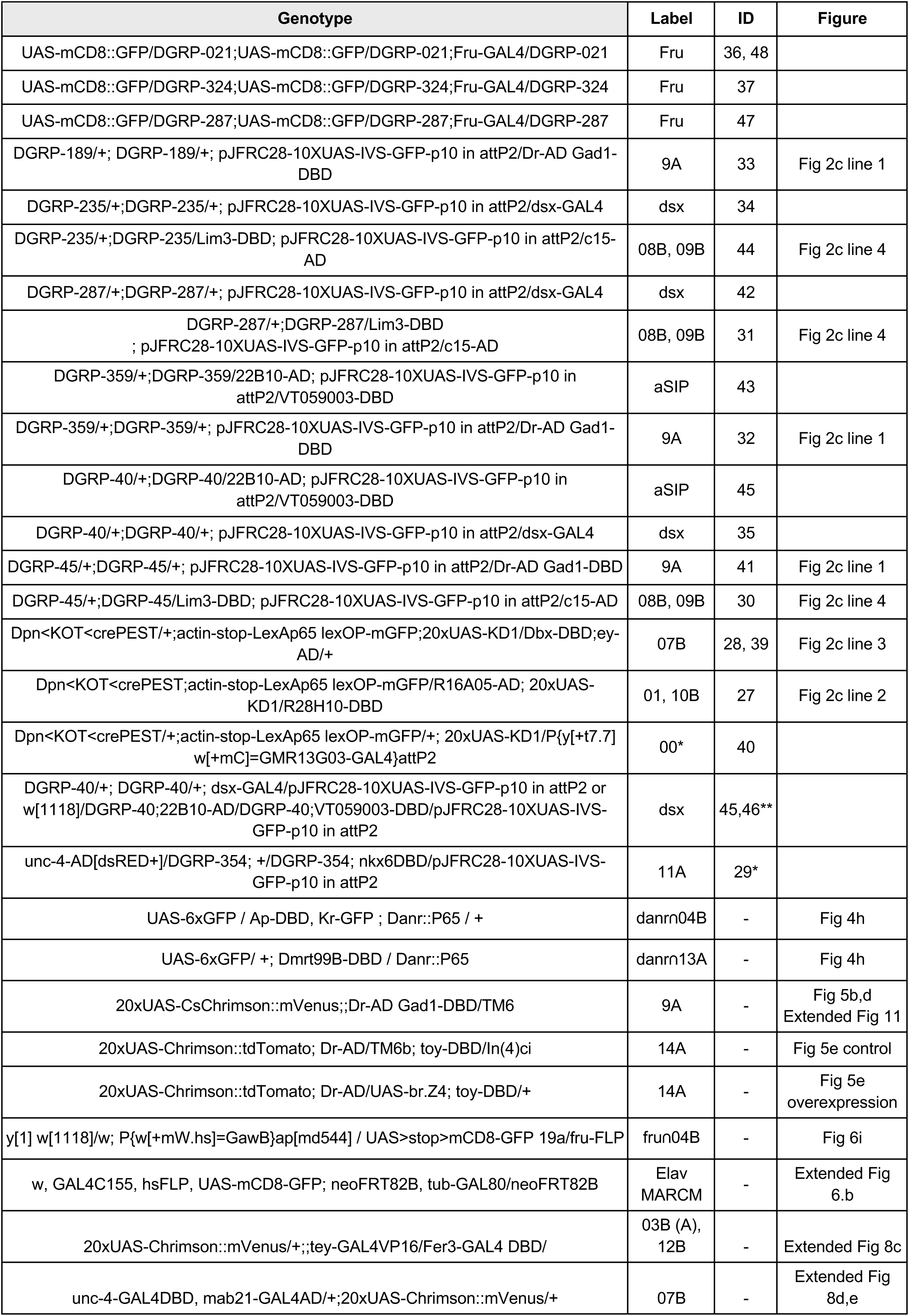
Genotypes used in this study. ID column corresponds to UniqueCrossID metadata column in the VNC seurat object. * line did not yield GFP expression. ** line was mixed at the dissociation step with that of genotype w[1118];22B10-AD;VT059003-DBD and was not used for any analysis depending on genotype identity.

**Table 3.**
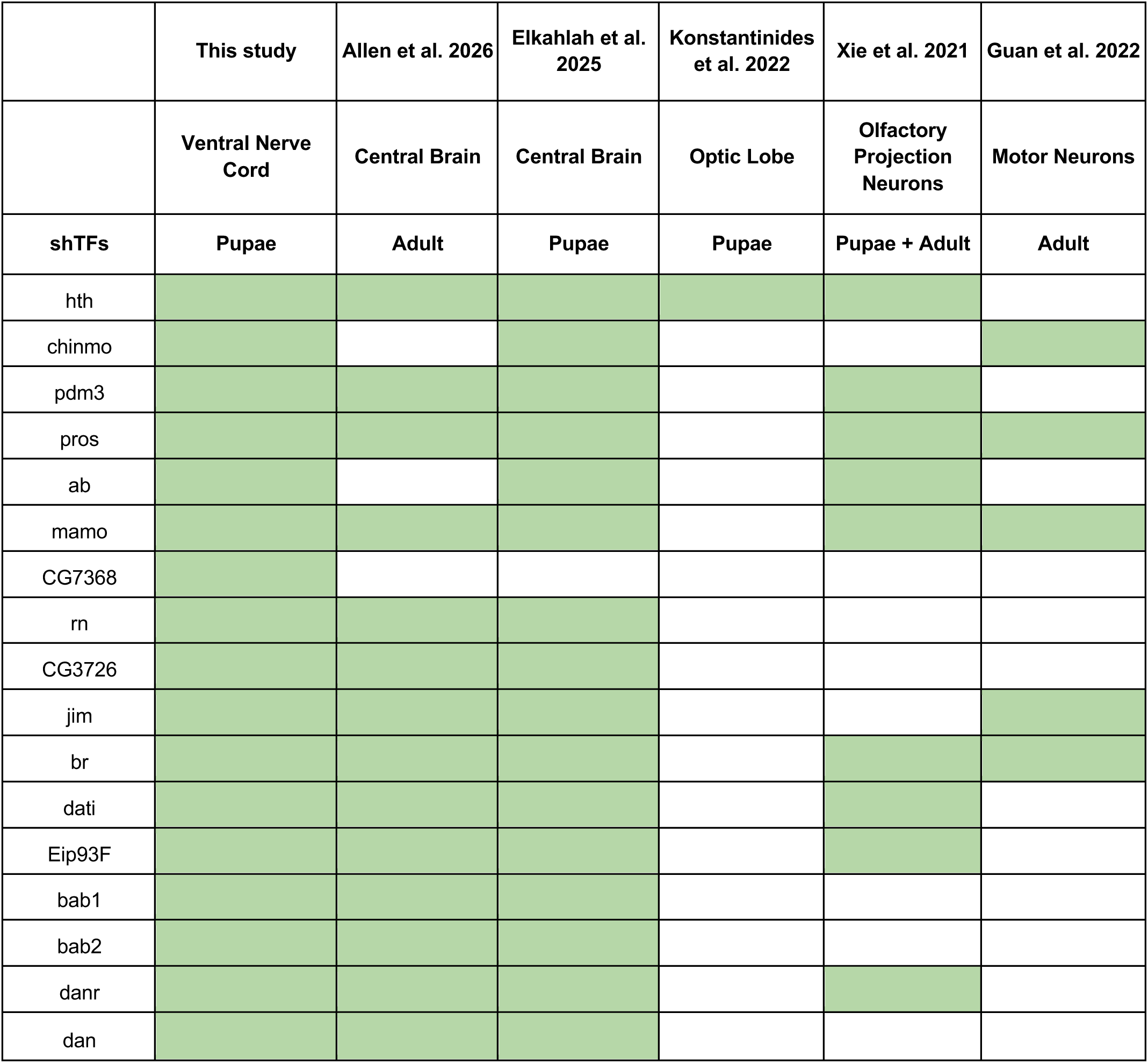
Summary of shTFs across studies. Columns correspond to different studies and rows report shTFs identified in this study.

#### Library preparation and sequencing

Samples were collected on 3 different days. Each day a total of 4 different cell suspensions (independent samples) were generated, each containing different combinations of genotypes, ages and sex. Single cell expression libraries were generated at the CRUK-CI sequencing facility (Cambridge, UK) using Chromium^TM^ Next GEM Single Cell 3′ HT v3.1. We aimed to load each sample in duplicate, 20,000 cells per lane. Before sequencing, libraries were inspected on a tapestation. Sequencing was performed on a NovaSeq6000 machine (Illumina) with the following sequencing parameters: 28 regular cycles of which 16 are 10x barcode and 12 are unique molecular index (UMI), 10 i7-index cycles, 10 i5-index cycles, 90 regular cycles.

### scRNAseq data processing

#### Cellranger

Data from the NovaSeq6000 sequencer was processed using CellRanger v8.0.0 using cellranger count function with default parameters. CellRanger reference index genome was built on genome assembly BDGP6.32^103^ supplemented with the coding sequences for the expected transgenes (Supplemental File 3). Before quality control, cell ranger total output was 805,970 cells. The median sample number of reads per cell was 39,142, the number of detected genes per cell 1,511 (min=183, max=8,543) and the median number of Unique Molecular Identifiers (UMI) per cell 4,893 (min=666, max = 847,868).

#### Demultiplexing

The output from cellranger was demultiplexed using applications from the Demuxafy docker container version 1.0.3 (https://demultiplexing-doublet-detecting-docs.readthedocs.io/en/latest/index.html). Single Nucleotide Polymorphisms (SNPs) information for all sequencing reads (Pileups) were generated using cellsnp-lite with the parameters “--chrom 2L,2R -p 22 --minMAF 0.1 --minCOUNT 100”. Out of the 4 fly chromosomes only chromosome 2 was used for demultiplexing because in our cross schemes it always comes from the DGRP lines, while the remaining chromosomes originate on the other stocks. The demultiplexing package Vireo ^102, 104^ was run twice on the pileups, once using a Variant Call Format (VCF) reference file and once without it (withVCF and noVCF in Extended Data Fig. 1). The VCF file for the first run was generated using the DGRP SNP data from the Aerts lab (https://resources.aertslab.org/DGRP2/NCSU/final/dm6/DGRP2.source_NCSU.dm6.final.SNPs_only.vcf.gz). The parameters used were “-N noSamples -p 4 --randSeed=1” where noSamples is the number of different genotypes present in the given sample. The noVCF run can identify all genotypes in an unbiased way but does not provide the identity of the groups, i.e. while it produces groups 1 to 7 it does not indicate which genotype each group corresponds to. Conversely, the run withVCF can identify most genotypes but is less efficient in identifying singlets due to the variable quality of VCF information across DGRP genotypes and the confounding effect introduced by DGRP heterozygosity of chromosome 2 (only one parent carried the DGRP chromosome). In order to use the demultiplexing results of the more efficient noVCF runs we used several source of information to assign groups to genotypes: (1) the correspondence between expected sex and roX1 expression, (2) expression of GFP transgenes, (3) comparison of the results from noVCF and withVCF Vireo results (Extended Data Fig. 1). In the rare cases where the genotype assignment provided by the withVCF run disagreed with the Sex or GFP transgene expression, these took precedence. Besides enabling assignment of metadata information such as sex and stage of the cell, demultiplexing also allowed removal of doublets, given that those cells have a high fraction of SNPs coming from different DGRP lines. Cells were considered doublets if they were classified as such in the Vireo noVCF run and excluded from further analysis.

#### Genes annotations

The list of transcription factors (TFs) comprises the union of genes annotated as TFs in flybase and TFs reported in Seroka et al 2022^105^. Annotation of cell surface molecules was taken from Seroka et al 2022^105^. Genes involved in cellular energy production and stress response correspond to the union of flybase gene lists:

> CYTOCHROME_C

> ELECTRON_TRANSFER_FLAVOPROTEINS,
>
> MITOCHONDRIAL_COMPLEX_III,
>
> MITOCHONDRIAL_COMPLEX_II,
>
> MITOCHONDRIAL_COMPLEX_IV_CYTOCHROME_C_OXIDASE_SUBUNITS,
>
> MITOCHONDRIAL_COMPLEX_I_SUBUNITS,
>
> MITOCHONDRIAL_COMPLEX_V_ATP_SYNTHASE_COMPLEX_SUBUNITS,
>
> heat_shock_prot

#### Quality Control

Cell ranger output was imported into an R object using sequentially the functions Read10X and CreateSeuratObject from the Seurat package version 4^106^ with default parameters. Some of the last analyses were done using Seurat version 5^107^. A first QC filter was applied to exclude cells with fewer than 700 features, more than 20% of mitochondrial genes and more than 50% of ribosomal genes. Genes expressed by fewer than 3 cells in the dataset were also excluded. Samples showed a bimodal distribution of UMIs and features across cells except for SITTD8 where the distribution was unimodal. For this reason we excluded SITTD8 from further analysis.

#### Data integration

Data integration was performed twice following the same pipeline (see below), to integrate the full dataset (including neurons and glia) as well as the subsets containing either neurons or glia.

Data integration was performed using the *rpca* algorithm in Seurat. Briefly, 3,000 features were found using SelectIntegrationFeatures, FindIntegrationAnchors was then run on those features using the 24h samples as reference and lastly, the anchors were used with the IntegrateData function with a k.weight = 100 (or number of cells -2 for small samples). This integration process removes variability due to stage and genotype. Cells with fewer than 3000 UMIs or in clusters with an average number of UMIs smaller than 3,000 were excluded from further analysis as these were deemed low quality. The resulting VNC dataset has a median of 23976.3 UMIs (min = 3,000, max = 565,460) and a median number of genes of 3,445 (min = 731, max = 8,428).

### Atlas Annotation

#### Neurons and Glia

To annotate neurons and glia we calculated, using the Seurat function AddModScores, enrichment scores for glial (*wrapper, repo, CIC-a, loco, CG10702, CG6126, Gs2, Egfr, Tret1-1, bdl, Zasp52, rols, ine, CG5404, CG14688, CG31663, ry, CG4752, betaTub97EF, CG32473, LManII, Eaat1, alrm, Eaat2, axo, Vmat, moody, Indy, zyd*), neuronal (*elav*,*nSyb)* or “other” (*CG11835, alphaTub85E, Act57B, betaTub60D, CG5080*) marker genes. Markers for the category “other” were chosen for being specifically expressed in a cell population lacking neuronal as well as glial markers. Gene ontology analysis identified several genes enriched in cells of mesenchymal origin among markers for the “other” category. Categorical assignment of cells to “Glia”, “Neuron” or “other” was done based on a cluster-based winner-takes-all criterion. The category “other” was excluded from any further analysis.

#### Glia types

Glial clusters were annotated according to the expression of previously reported markers ^108^. Specifically *sim* expression was used for midline glia, *CG6126* for MAP, *ltl* for subperineurial, *alrm* and *Tre1* for astrocytes, *wrapper* for cortical and *axo* for ensheathing glia. Similarly to neurons and glia, glial identity was assigned based on a cluster-based winner-takes-all criterion.

#### Neurogenesis wave and Maturation State

The adult brain contains primary neurons, born in the embryonic phase, and secondary neurons, born during larval stages. To identify these distinct populations in our atlas we used the markers Imp (primary) and dati (secondary) ^36–38^. First we assigned categorical identity to neurons expressing high levels of markers (primary: Imp > 2, secondary: dati > 2 in the stage-integrated data slot), producing nearly completely mutually exclusive labels. We reasoned that neurogenesis wave information might be more prominent in earlier developmental stages and used the obtained labels to train a classifier using the devCellPy machine learning framework ^109^ with parameter rejCutoff = 0.5 and testSplit = 0.1. The trained model was used to predict primary and secondary labels across the atlas. Expression data for training and classification included all 3000 genes from the stage-integrated data slot.

The stem region of our atlas contains cells with low expression of *dati* and markers of mature neurons, e.g. *brp* and *nSyb*, and it is devoid of 36h and 48h cells, suggesting it corresponds to neuronal precursors. We annotated these neurons using a similar strategy as above, but using *brp* (> 1.5) for mature neurons and the chromatin remodelling factor *Phs* (> 1.5) for immature ones. Similarly to neurogenesis wave we trained the classifier on 6h data and then used the model to predict labels across the atlas. In both cases training and prediction was performed 50 times starting from the same training set and the consensus result used for the final assignment.

#### Cell Cycle Phase

Cell cycle annotation was done using the Seurat function CellCycleScoring. Marker genes for the different phases of cell cycle were obtained from Tirosh et al. 2016 ^110^.

#### Neurotransmitter identity

Each neuron in the atlas was assigned a fast neurotransmitter identity based on the highest expression level among the markers VAChT (cholinergic), Gad1 (GABAergic), and VGlut (glutamatergic). Neurons were assigned a monoamine class (tyraminergic, octopaminergic, dopaminergic, serotonergic and histaminergic) if they co-expressed the vesicular monoamine transporter (Vmat) and monoamine specific markers (tyramine = Tdc2+, Tbh-, octopamine = Tdc2, Tbh; dopamine = Ddc, ple; serotonin = Trh, Ddc; histamine = Hdc).

#### Hemilineage

To provide experimental evidence for hemilineage assignments we included among the sequenced samples transgenic lines where GFP expression is specifically driven in lineages/hemilineages (09A, 01+10B, 07B, 08B+09B). This was obtained either through split-GAL4 intersectional approach (09A - line 1: Dr-AD/Gad1-DBD; 08B + a subset of neurons from 09B - line 4: Lim3-DBD/c15-AD)^43,111^ or through an immortalisation-based strategy (01+10B - line 2: R16A05-AD;R28H10-DBD, 07B - line 3: Dbx-DBD/ey-AD) ^39,112^.

Annotation of hemilineages was performed using a stepwise approach, integrating information from multiple published and unpublished sources (summarised in Supplemental File 1). We first assigned hemilineages with strong supporting evidence from previous studies, which helped reduce the number of unassigned “orphan” hemilineages requiring further annotation. We started by mapping the adult VNC atlas ^31^ onto the neuronal subset of the developmental dataset and transferring the existing hemilineage annotations to the pupal dataset using the Seurat functions FindTransferAnchors and TransferData with 3000 variable features and 200 principal components. Then, selectively in secondary neurons, we expanded adult-derived annotation by taking an iterative cluster-based approach. Hemilineage labels were expanded to all cells within a cluster if more than 50% of its neurons were already annotated with the same hemilineage. This procedure was performed iteratively, from high clustering resolution, i.e. small clusters, to low clustering resolution, i.e. big clusters. Annotations were then discarded if, after the iterative procedure, they accounted for less than 20% of a cluster at an intermediate clustering resolution (resolution = 1.2). Finally, we applied a winner-takes-all strategy when the annotation represented at least 30% of the cluster at a high clustering resolution (resolution = 20). We then resolved ambiguous hemilineage identities in cases where the adult annotation grouped two hemilineages together, but our dataset showed them as clearly separated clusters (e.g. 05B/06B). We calculated differentially expressed genes (DEGs) between the two ambiguous clusters using FindMarkers, which confirmed cluster-specific genes and assigned cluster identity according to the relative expected hemilineage size in MANC^113^. For most hemilineages, identity was further corroborated by previously published split-GAL4 lines with experimentally validated expression patterns (see Supplemental File 1)^114^. Next we annotated hemilineages not covered in the Allen et. adult dataset, but expected to contain more than 10 secondary neurons based on the MANC annotation. For this we combined evidence from neurotransmitter gene expression, GFP expression from labelled lines, markers from the literature, and comparison between cluster size and number of neurons expected by EM. Iteratively we re-clustered only neurons without hemilineage annotation and assigned them to a specific hemilineage based on the expression of identified new markers. For a detailed hemilineage-by-hemilineage summary of the annotation strategy employed refer to Supplemental File 1. This led to an initial assignment of 48h secondary neurons. Annotations were discarded if they accounted for less than 30% of a cluster at an intermediate clustering resolution (resolution = 1.2). This led to the identification of high-confidence assignments (Fig. 2c) in 48h secondary neurons that were used to train a classifier using the devCellPy machine learning framework ^109^ with parameter rejCutoff = 0.5 and testSplit = 0.1. The trained model was used to predict hemilineage labels across the atlas at all stages and expanding to primary neurons, using rejCutoff = 0.3 for prediction. Expression data for training and classification included all 3000 genes from the stage-integrated data slot. Training and prediction was performed 50 times starting from the same training set and the consensus result used for the final assignment.

Note that since secondary neurons of hemilineage 15B are exclusively motor neurons, lineage propagation to primary neurons resulted in the broad assignment of 15B identity to all motor neurons, which share many molecular features. Primary 15B neurons are to be intended more broadly motor neurons.

Subesophageal zone specific hemilineage 27X and 03A in LB (labium, glutamatergic unlike VNC homologues) were assigned based on expression of eya^39^ and fd96Cb respectively. fd96Cb (CG11922) is associated with the GMR line R74D11 that drives specific expression in one hemilineage in LB ^115^.

#### Soma Neuromere

We annotated soma neuromeres based on Hox genes known to be expressed differentially along the VNC anterior-posterior axis (Dfd, Scr, Antp, Ubx, abd-A and Abd-B). Dfd+ and Scr+ cells belong to hemilineages originating from the gnathal ganglia (labial/LB, maxillary/MX and mandibular/MD)^46^. Their presence at 6h stage in our dataset likely reflects imprecise excision of the VNC from the brain at this early time point when the neck constriction has yet not formed (Extended Data Fig. 6). We assigned all Dfd+ or Scr+ cells to GNG^46^. In those cases where we had information about hemilineage-specific expression of Dfd and Scr in the different GNGs, we refined annotation (i.e. 03B). A more detailed annotation of the GNG segments will be covered in a different manuscript (*in prep.*). All abd-A+ or Abd-B+ cells were annotated as A regardless of the developmental stage given that those markers are not expressed outside the abdominal segments. For the annotation of T1, T2, T3 and abdominal cells not expressing abd-A or Abd-B, instead, we first annotated cells at 48h and then transferred annotation to earlier time points, due to significant variation of *Antp* expression across pupal stages. Since the expression pattern of Antp and Ubx is hemilineage-specific, we treated each hemilineage independently. In cases where immunostaining data from L3 larvae indicated that expression of *Antp* and *Ubx* genes deviated from the canonical model, we used these data as guidance. A summary of Hox genes expression derived by these data is reported in Supplemental File 4; specific images are available upon request. The workflow followed to annotate soma neuromeres is shown in Fig. 3b. For each hemilineage we generated violin plots showing expression by cluster at a high clustering resolution for *Antp*, *Ubx* and *abd-A*. These plots helped choose expression thresholds to assign cells to T1, T2, T3 or A, a critical task when the difference between the different segments is dictated by the level of expression and not by the quality of the gene expressed. Thresholds used for each hemilineage are summaries in Supplemental File 4. Thresholding based annotation was used as input for further annotation using devCellPy. Cells were excluded from the training set if their assigned label represented less than 10% of the total cells in their cluster. To prevent overtraining, only half of the labelled cells were used to train the classifier. Multi-layered classification provides hemilineage-specific soma neuromere models which are essential to capture the hemilineage-specific Hox gene expression differences. Stage integrated expression values were used to account for expression differences due to developmental stage and allowed transfer of the model learnt using 48h data to earlier stages (as done to assign hemilineage identity). The training parameters used were: rejCutoff = 0.5, testSplit = 0.1. The obtained classifier was used to predict the full dataset with prediction parameters: rejCutoff = 0.3. Training and prediction were run 50 times starting from the same training set. Results were collated and used to obtain a consensus result used for final assignment.

We used the FindAllMarkers function to identify segment specific markers at 48h for hemilineages with complex UMAP trajectories. The 10 positive markers with lowest p-value, expressed in at least 80% of the cells in the segment are shown in Extended Data Fig 3.2a.

#### Fruitless clones in the atlas

To identify neurons belonging to fruitless clones in the atlas, we first selected fru-expressing cells using a gene expression threshold of 0.7. Next, we clustered all hemilineage–segment trajectories using the 17 shared transcription factors (shTFs; see Shared Peaks Detection) as input dimensions. The clustering resolution was adjusted to yield a number of clusters approximately equal to half the number of neuron types expected based on MANC annotation. Clusters were considered fru+ if at least 50% of cells were. Cells in fru+ clusters were assigned to clone identity based on matching hemilineage and segment (see Identification of neurons belonging to fru MARCM clones in the connectome for more explanations about fru clones).

### Primary - Secondary analysis

For the analysis of primary and secondary neurons we split the dataset as follows: cells annotated as primary > primary; cells annotated as secondary or early secondary > secondary. 2000 variable features and 200 PCs were re-calculated for each subset independently in the integrated assay and used to generate the UMAP embedding shown in Fig. 1d (separated plots), using default parameters for the appropriate Seurat functions. For cluster-size comparison to EM types and inter-cluster overlap analysis we excluded 6h cells, as these were often separated to the rest of the stages, likely due to the presence of subesophageal zone cells, cells committed to programmed cell death in the abdomen and their low degree of maturation; this incomplete integration would degrade the power of the analysis, e.g. resulting in 6h-only clusters. The atlas coverage after removal of 6h cells is 28x. We re-calculated UMAP embeddings and nearest neighbours (k=20) using the same 200 PCs, reasoning that the information carried by the 6h cells could help better separation of clusters. We then calculated clusters at different clustering resolutions, using Seurat FindCluster function with default parameters, and chose resolution 16 for the analysis in Fig. 1h-i.

For the constellation plots, we first computed the number of edges between each pair of clusters based on the nearest-neighbor graph (integrated_nn). Self-edges (within-cluster connections) were excluded. In cases where multiple cells in A had the same nearest neighbour in B only one of those connections was kept and the rest excluded. For each directed edge (from cluster A to B), the raw edge count was normalized by the number of cells in the target cluster (B), resulting in a directional normalized weight. To take into consideration the bidirectional comparison, we averaged the two reciprocal normalized weights between each cluster pair (A→B and B→A). This final edge weight thus reflects the average fraction of inter-cluster nearest neighbours. For the graphs, we retained only edges with a weight greater than 5%. Clusters without inter-clusters edges after filtering were labelled in brown. For the density distribution analysis, no filtering was applied and, for each cluster, the strongest overlapping edge was retained (a measure for the distance from the closest cluster). Statistical significance between distributions was assessed using the Wilcoxon rank-sum test.

To compare the number of cells per cluster to the number of neurons per type in the connectome we divided the cluster size by the expected coverage, thus making the two numbers comparable. For primary neurons in MANC we retained only neurons labelled as “primary”, while for secondary neurons we retained only “secondary”. Early secondary neurons were excluded from the analysis because of the uncertainty in assigning them with confidence to the primary or secondary transcriptional group (we believe that most primary neurons in the atlas correspond to primary+early secondary in MANC, where birthtime annotation of these neurons has low confidence). Neurons sharing the same “serial” annotation were considered a single type, while those lacking a “serial” annotation were grouped by their “type” label. The number of neurons in each serial/type group was then divided by two to account for the fact that coverage was calculated on the hemi-connectome.

### Trajectories analysis

#### Pseudotimes and differentially expressed genes

For trajectory inference, UMAP embeddings were recalculated for each hemilineage using Seurat. Trajectory analysis was performed using Monocle3^54–56,116^ and cells falling outside the main trajectory or in regions lacking 48h cells removed using the monocle3 function choose_cells. For discontinuous trajectories (i.e. trajectories defined over more than one partition), gaps in pseudotimes between partitions were removed. Trajectories of secondary neurons were analyzed independently for each hemilineage–segment combination. Hemilineage 15B was excluded due to the absence of a clearly defined trajectory, and 08B-T3 omitted due to its complex and discontinuous structure. Genes differentially expressed along trajectories were calculated using the monocle3 function graph_test and expression data from the stage-integrated slot, comprising all stages.

#### Trajectories alignment

To align trajectories we used G2G ^59^, which performs gene-level trajectories alignment. All trajectories were compared to the reference (03A-T1, chosen as reference for being one of the longest in the dataset) using a fixed number of bins (25). We ran G2G twice with two different gene sets. For the initial alignment we used a set of 40 TFs, corresponding to differentially expressed TFs along trajectories (q_value < 0.01, Moran’s index > 0.3) shared by at least 50% of the hemilineages, after taking the union across segments to limit the effect of short trajectories that would result in the exclusion of genes expressed at high pseudotimes values (ab, Antp, bab1, bab2, br, CG14431, CG3726, CG7368, CG9932, chinmo, chn, crol, dan, danr, dati, Eip93F, fru, hang, HmgZ, hth, jim, klu, l(3)neo38, luna, mam, mamo, NK7.1, noc, nvy, Octbeta2R, pdm3, pros, rn, scrt, tio, tna, tsh, Ubx, zfh1, zld). For the second alignment we used only 17 shTFs (see below).

To warp pseudotimes to the reference we fitted a linear model to the G2G output, matching bins between query and reference trajectories.

Combinations with fewer than 30% of the mean cell count across trajectories (i.e. 09B–T1, 17A–T1, 09B–T3) were excluded from warping and peak conservation analysis.

#### Shared peaks detection

For each trajectory, expression peaks were identified using the find_peaks function from the ggpmisc package, applied to expression profiles generated with a modified version of Monocle3 get_fitted_values function. Peaks were calculated for each of the 40 TFs, provided they were differentially expressed along the corresponding trajectory. Similarly, we calculated expression peaks for the average expression profile obtained by averaging expression in bins after warping trajectories to the reference space (average peaks). Average expression included only trajectories for which the given TF was differentially expressed.

For peaks matching, each reference peak was warped to the reference space and assigned the identity of the closest peak in pseudotime. A peak was considered shared if it lay within 5 pseudotime units and was at least half the height of the corresponding average peak. Genes were retained if at least one peak was shared in 50% or more of the analyzed trajectories. This procedure identified 17 shTFs. For each trajectory, we ranked the positions of all trajectory-specific peaks for shTFs and computed a consensus order across trajectories using the consrank function from the ConsRank package. Trajectories with fewer than 20% of the total number of peaks were excluded from the ranking analysis (19A-T1, 22A-T1, 03B-T3).

### Central brain dataset analysis

Data for the SLPad1 trajectory shown in Fig. 5 are derived from a corresponding developmental transcriptional atlas of the brain (without optic lobes), which will be described in detail in a separate manuscript.

Briefly, data collection was performed as described for the VNC, with few differences. 1- For the brain we collected samples at 6, 24, or 48h. 2- Up to 8 samples per genotype were pulled in the same tube prior to dissociation and a total number of 32 brains were pulled. 3 - Ten independent experiments were submitted to the CRUK-CI facility for single-cell RNA-seq using 10x Genomics Chromium Next GEM Single Cell 3′ kits (v3 or v3.1). Each sample was loaded on one or more lanes with 10,000 cells per lane.

Downstream analysis was carried out as described for the VNC. Although a complete hemilineage annotation is not yet available for the brain, we identified a trajectory of secondary neurons marked by Lim3 expression, with a fru-expressing cluster located opposite the Imp-positive tip - likely corresponding to the aSP-f clone. Cells along this trajectory (Lim3+/nompB+ cells and Lim3+/Imp+ cells) were manually selected in the stage-integrated UMAP embedding, which facilitated identification of the best-matching clusters. The manually selected cells were matched to a single cluster in the stage-integrated space (83 at resolution 1.2 using the Louvain algorithm). Cells labelled as precursors were excluded from the final SLPad1 object. Data for subesophageal zone trajectories (03A and 03B) comprised 6h data from the VNC atlas. Cells along these trajectories were manually selected from stage-integrated UMAP embedding, to ensure inclusion of all cells. Trajectory analysis was performed as previously described for the VNC.

To calculate the expression correlation between the shTFs in the reference 03A-T1 and all hemilineages in the VNC and the 4 hemilineages from the brain we followed:

1. Expression values were extracted from the data slot.
2. Expression was pooled in bins of 0.5 width of warped pseudotime units (i.e. pseudotime values after registration to reference).
3. A gaussian smoothing was applied using the smoother package function smth.gaussian with parameters: window = 0.2, alpha = 0.1 and tails = TRUE.
4. The correlation between each pair of shTFs from the reference and the hemilineage under analysis was calculated using the cor function from the stats package and the parameter: use = “pairwise.complete.obs”.
5. A single value for each hemilineage under analysis was calculated as the mean of the 17 correlation values.
6. To calculate the shuffled values, the process was repeated 200 times for each comparison while the identities of the shTFs in the reference were shuffled.

### Sex differences in the transcriptional atlas

#### Differential abundance analysis

Numerical differences between sexes were analysed using miloR^67^, a cluster free method well suited to identify local differences in continuous hemilineage trajectories. We followed the pipeline suggested by developers. Stages were analysed separately and stage-integrated data were used to compute the shared nearest neighbour graph. We kept default function parameters with the exception of: k = 10, d = 200 for buildFromAdjacency and prop = 0.2 for makeNhoods.

#### Detection of sexual dimorphisms

We reasoned that sexually dimorphic neurons should be characterised by an increased molecular distance compared to isomorphic neurons. To quantitatively describe such distance at the single-cell level, we defined a correction score value as the sum of the differences of the per-gene expression level before and after sex-driven variability regression using the rpca integration algorithm in Seurat (similarly to what was done for stage integration). Integration was performed independently for each hemilineage-segment combination on cells from all stages. To calculate correction scores in females, male samples were kept as reference, and vice-versa. To calculate statistically significant differences between scores at 24 and 48h, cells were clustered using the 17 shTFs as input dimensions as described for fru+ clones annotation.

The same clusters were used to calculate DEGs between males and females at the different stages using Seurat’s FindMarkers function with default parameters (considered significant if p_val_adj < 0.05).

### Immunostainings and confocal Imaging

Unless stated otherwise, immunohistochemistry was done as described^117^.

Primary antibodies: mouse anti-nc82 (DSHB, AB_2314866) 1:40, chicken anti-GFP (Abcam, ab13970) 1:1000, guinea pig anti-Hth 1:500, rat anti-Dan 1:500, mouse anti-Br 1:50, rabbit anti-Eip93F 1:2500,

guinea pig anti-Mamo 1:1000, rabbit anti-Bab1 1:1000, rat anti-Bab2 1:1000, rat anti-Pdm3 1:500. Secondary antibodies: Alexa-568 anti-mouse (Invitrogen) 1:400, Alexa-488 anti-chicken (Invitrogen) 1:400. Specimens were mounted in Vectashield (Vector Labs) on positively charged slides.

Confocal stacks were acquired using a Zeiss LSM 880 microscope with Airyscan and motorised stage at 768 × 768 pixel resolution every 1 μm (0.46 × 0.46 × 1 μm) using Zeiss EC Plan-Neofluar 40×/1.30NA oil objective or LD LCI Plan-Apochromat 25x/0.8NA multi-immersion objective. All images were acquired at 16-bit color depth. Maximum projections of z stacks were made in Fiji.

#### Hox genes expression analysis in L3 larvae

Sparse lineage labelling was obtained through the generation of MARCM (mosaic analysis with a repressible cell marker)^118^ clones. Briefly, flies of the appropriate genotypes were crossed and females allowed to lay on grape juice agar plates for 12 hours at 25°C. After incubating at 25°C for 12 hours, the embryos were heat-shocked in a 37.5°C water bath for 30 minutes, rested at room temperature for 30 minutes, then heat-shocked again for 45 minutes. Larvae were reared on 4-24 instant fly media (Carolina Biological Supply) at 29°C to increase expression of the GAL4^C155^ driver and visibility of MARCM clones. The w, GAL4C155, hsFLP, UAS-mCD8::GFP; FRT82B; tubP-GAL80 stock was generously shared by Jay Parrish (University of Washington, Seattle) and the P{ry[+t7.2]=neoFRT}82B ry[506] stock obtained from the Bloomington Drosophila Stock Center (Indiana University).

Nervous systems were dissected from wandering third instar larvae of both sexes in PBS (pH 7.2), fixed in 3.7% buffered formaldehyde at room temperature, then washed in 0.3% PBS-TX (PBS with 0.3% Triton-X100). Fixed samples were blocked in 2% normal donkey serum (Jackson Immunoresearch Laboratories, West Grove, PA, USA) in PBS-TX for 30 min, then incubated for several days at 4°C with primary antibodies: rat anti-mCD8 (Invitrogen, CALTAG, cat. no. RM2200) at 1:100, mouse monoclonal anti-neurotactin (BP106) at 1:30 or mouse monoclonal anti-Antp (8C11) at 1:500 (Developmental Studies Hybridoma Bank, University of Iowa, Iowa City, IA, USA), and rabbit anti-Ubx (7701) at 1:1000^45^. After primary antibodies were washed out in PBS-TX at room temperature, tissues were incubated overnight at 4°C in a 1:300 dilution of Alexa 488-conjugated donkey anti-rat IgG, Alexa 555-conjugated donkey anti-mouse IgG, and/or Alexa 647-conjugated donkey anti-rabbit IgG (Invitrogen). Following additional washes in PBS-TX at room temperature, tissues were arranged on polylysine-coated coverslips, cleared in xylene, and mounted in DPX (Sigma-Aldrich, St. Louis, MO, USA).

Slides were imaged using a 40x oil objective on a Leica SP5 Spectral Systems confocal microscope. Z-stacks were collected sequentially with averaging at 0.5 to 1.0 μm intervals. Raw data stacks were imported into Fiji (https://fiji.sc/) and either merged or projected into 3D representations for lineage identification based on morphology, neuroblast location, projections into a landmark neurotactin scaffold, and/or Ubx expression, using our published atlases^40,44,45^. Confocal stacks were processed and assembled into figures using Fiji and Affinity Publisher. Images are shown as maximally projected 2D views of subsetted confocal stacks, without removing potentially obscuring clones that can be resolved in 3D representations.

#### Validation of segments annotation

VNCs were dissected and fixed as previously described^43^. Primary antibody staining was performed sequentially: samples were incubated overnight at 4°C with mouse anti-nc82 (1:25, DSHB) and either preabsorbed chicken anti-GFP (1:1000, Life Technologies) or rabbit anti-GFP (1:1000, Life Technologies), followed by an overnight incubation with rabbit anti-tey^119^ (1:200, gift from A. Stathopoulos) or guinea pig anti-kn^120^ (1:200, gift from W. Moore), respectively. Corresponding secondary antibodies (Alexa-568 anti-mouse, Alexa-488 anti-chicken or Alexa-488 anti-rabbit, Alexa-633 anti-rabbit or Alexa-647 anti-guinea pig, 1:500, Life Technologies) were applied overnight at 4°C. Samples were washed in PBS plus 0.1% Triton X-100, positioned onto lysine-coated coverslips and dehydrated through an ethanol series, cleared in xylene and mounted in DPX^44^. Imaging was performed using a Nikon ECLIPSE Ti2 confocal microscope equipped with a Nikon Plan Apo 40× oil-immersion objective. Confocal stacks were acquired at 1024 × 1024 resolution with sequential scanning and z-steps of 0.5–1.0 µm.

#### EdU labelling

EdU labelling experiments were performed as described in Daul et al. 2010^121^. Briefly, newly hatched larvae were collected for 1 hour, transferred to fresh food and raised at 25°C until the appropriate stage for the EdU pulse. Larvae were then transferred to EdU-conditioned food and allowed to eat for 7 to 8 hours. After the pulse, larvae were transferred to fresh unconditioned food till eclosure. Adult brains and VNCs were dissected, fixed and stained. EdU detection with Click-iT™ EdU Imaging Kit with Alexa Fluor™ 647 (Thermo Fisher) was performed following manufacturer’s instructions and the following immunostaining was performed as previously described^117^. Primary antibodies: mouse anti-Br-core monoclonal (1:50) and rabbit anti-Bab1 polyclonal (1:1000). Secondary antibodies: Alexa-594 goat anti-mouse-IgG1 (1:1000) and Alexa-594 goat anti-rabbit (1:1000). Confocal images were acquired using a Leica TCS SP8 3X gated STED confocal microscope equipped with a white light laser using optimal excitation frequencies for each fluorophore and a 40x/1.1NA Water objective. Care was taken to acquire channels independently to avoid bleedthrough between 594 and 647 fluorophores. Confocal stacks tiling the VNC and brain were stitched either by the microscope software with default parameters or using the pairwise stitching plugin in Fiji with default parameters ^122^. Fluorescent signals for Br, Bab1 and EdU were segmented using channel-specific Ilastik models (Ilastik 1.4.1^123^) trained on multiple images. Simple Segmentation outputs were generated from the Pixels Classification pipeline and transformed into binary masks in Fiji. Expression overlap was quantified as the number of pixels in the confocal stack positive for both shTF and EdU, divided by the total number of shTF pixels.

### Connectomics data analysis

Summary statistics refers to MANC v1.2.3 (root node: “1ec355123bf94e588557a4568d26d258”). Data are available from https://neuprint.janelia.org. The total number of annotated intrinsic neurons (i.e. neurons with soma in the VNC volume, corresponding to classes ascending neuron, efferent ascending, efferent neuron, intrinsic neuron, motor neuron) is 15,760. Throughout the analysis numbers refer to one side, under the assumption that the VNC is bilaterally symmetric and that the same number of neurons is expected on each side. For example, to calculate aggregate coverage: 302,765 / 7,880 = 38.4.

For the analysis and visualisation of connectomics data we used packages from the natverse toolbox ^124^. Dataset-specific packages for the male VNC (MANC, https://github.com/natverse/malevnc), the female VNC (FANC, https://github.com/flyconnectome/fancr) and the female brain (fafb/flywire, https://github.com/natverse/fafbseg).

#### Similarity to nearest neighbour

We downloaded skeletons for all neurons in the MANC dataset, created dotprop representations and used them to run an NBLAST^125^ comparison. The scores for the best matches for each primary and secondary neuron were used to create the morphological comparison. Similarly, we downloaded the connectivity information for all neurons in the MANC dataset and used it to create cosine similarity comparisons within neurons from the left side and within neurons from the right side. The scores for the top matches for each primary and secondary neurons were then used to create the connectivity comparison. The statistical significance of observed differences was evaluated using a Wilcoxon rank-sum test.

#### Identification of 14A types matching light-level stainings

To identify the neurons shown in Fig. 5f, we queried Clio for all neurons annotated as hemilineage 14A, soma_side = RHS, soma_neuromere = T1, and birthtime = secondary or early secondary. We manually excluded neurons whose projections did not match the morphology observed in light-level images, yielding 17 seed neurons (IDs: 17237, 21776, 23665, 23829, 24397, 25482, 25805, 29164, 31376, 32854, 34040, 42121, 43194, 46063, 103686, 155414, 166875). All neurons sharing the same systematic_type annotations as these seeds were selected to generate the set shown in Fig. 5f (left). We then identified additional neurons that follow the same axonal bundle as the seed set (reduced in Br OE condition). Finally, the remaining 14A, RHS, T1 secondary/early-secondary neurons were used as seeds to select contralateral neurons and neurons in other neuromeres with matching systematic_type annotations, generating the dataset shown in Fig. 5f (right).

#### Male-Female connectomes matching

The lack of extensive neuronal typing of the female EM dataset (FANC) or matching to homologous male neurons in MANC, motivated us to develop an algorithm that uses a two stage pipeline to match single neurons from one datasets to one in the other. This allows the study of homologous neurons, the identification of potentially dimorphic neurons, with low score or missing altogether, and the transfer of cell type labels from MANC to FANC.

In the first stage of the process, a matching algorithm assigns neurons maximising morphology similarity (metric: NBLAST) across the dataset. In the second stage, a new assignment round, constrained by morphology similarity from the first round, aims to maximise connectivity similarity (metric: cosine similarity).

Assignment based on morphology > NBLAST matches

1. Neuronal skeletons were downloaded for both MANC and FANC datasets;
2. FANC skeletons were transformed into MANC space^126^;
3. Skeletons were converted to Dotprops.
4. Morphological similarity was calculated between neurons from each dataset as the mean normalised NBLAST score.
5. An implementation of the Hungarian algorithm calculated NBLAST matches between MANC and FANC neurons as those that maximised the global similarity between datasets, as measured by NBLAST scores from the assigned neurons.

Assignments by connectivity > connectivity matches

This assignment requires prior knowledge of neuronal identity, i.e. to evaluate how similar the connectivity of one neuron in MANC is to a neuron in FANC the identity of their partners, at least partially, must be known.

1. We established an initial panel of 50 homologous neurons between MANC and FANC to be used to calculate cosine scores between neurons from both datasets. These included highly similar neurons with high NBLAST score) and a single good match, thus bona fide homologues. We then selected the 2,000 neurons with the greatest difference between the top two NBLAST scores, a proxy for match uniqueness. From these,we selected the 1,000 neurons with the highest number of synapses,with the rationale that a large synapse count correlates with large neuronal size, and this, in turn, is more likely in primary neurons, which tend to be unique and have single matches across datasets. Neurons having many synapses are more useful to probe connectivity, as they connect to the largest possible fraction of un-labelled neurons. From these 1,000 we randomly selected a subset of 50.
2. In each dataset we identified the 500 neurons most connected to the panel of 50;.
3. We then computed cosine similarity between MANC and FANC, within these panels of 500 neurons, considering only connections to the panel of 50 neurons.
4. We matched MANC to FANC neurons based on the cosine scores, but only retained matches with a high NBLAST score (as a fraction of the top NBLAST match, e.g. if a neuron’s top match had a score of 0.8 then only neurons with NBLAST scores >0.8 * threshold would be considered) to ensure both high connectivity and morphological similarity. We therefore transferred homology labels from MANC to FANC, increasing the panel of labelled neurons from 50 to 550.
5. We repeated steps 2, 3 and 4 until all neurons in the dataset were analysed.
6. The NBLAST threshold used in 4 was progressively lowered to include additional match candidates until a predefined minimum threshold was reached. For the table below thresholds used went 0.95 -> 0.65 in 0.1 decrements.
7. The best one-to-one matches were recorded as the final “connectivity matches”, and the full history of match assignments was noted.

When compared with manual assignments between FANC and MANC recently published for ascending neurons^20^, the automatic assignment algorithm finds matches that in 85% of the cases are in agreement with the manual type assignment.

The results of the matching can be found in Supplemental File 5.

#### Hemilineage annotation of FANC neurons

To identify all neurons belonging to selected lineages in FANC we started from best matches from the MANC-FANC automatic matching results (same match for NBLAST and connectivity AND connectivity match score > 0.3). This allowed us to identify seed planes in the FANC EM volume containing most or all of the selected IDs from each hemilineage (typically the soma tract, or a point of entry to, or exit from, the VNC). Any extra neuron in the seed plane that appeared to belong to the bundle formed by the neurons in the hemilineage was marked as a potential candidate to belong to the hemilineage. Any obvious outlier was marked as an outlier and any neuron obviously belonging to other hemilineages was marked as “other”. Any neuron whose morphology was not clear due to a necessity for proofreading (e.g. loose soma, massive glia) was added to the list of neurons to be proofread, to ensure correct inclusion/exclusion from the hemilineage of interest. As one seed plane might not contain all of the neurons of a given hemilineage, extra seed planes were considered. All newly found neurons were coarsely proofread. Upon proofreading, neurons were re-matched to MANC and the final list of potential hemilineage markers was manually curated to obtain a final annotation, reported in Supplemental File 6.

#### Identification of neurons belonging to fru MARCM clones in the connectome

Previously published confocal stacks of segmented fru+ MARCM clones in VNCIS1 reference space^69^ were transformed to MANC space using functions from the navis package. Briefly, .nrrd files were read with navis.read_nrrd, thresholds were identified using np.quantile for the 99% and the images were then converted to meshes using the function navis.mesh and the calculated threshold. The new meshes in VNCIS1 space were processed using navis.xform_brain with parameters source VNCIS1 and target MANC. The resulting meshes were converted to neuroglancer precomputed objects using navis.write_precomputed. These were used to implement a neuroglancer (https://neuroglancer-docs.web.app/) layer that could be visualised in MANC space.

We identified MANC neurons potentially belonging to fruitless clones on the left hemisphere. For each clone, we first determined its lineage by loading neurons from candidate lineages into neuroglancer alongside the corresponding clone mesh. We looked for overlap of soma tracts. Once the lineage had been identified, we loaded all neurons from one hemilineage at a time and discarded those with processes outside the mesh. The remaining neurons were considered best candidates (*seed neurons*). When possible, we tried to identify as many *seed neurons* as predicted by light-level clones. To refine the selection we took a by-type approach: neurons were retained if at least 50% of the neurons in type were included in the seed (exception: Tr flexor MN in dPr-e, as this type comprises neurons from different lineages). When this procedure retrieved neurons with processes clearly outside the mesh, the type was manually excluded. When the number of neurons retrieved largely exceeded the expected number, EM meshes for individual types were downloaded, converted to VNCS1 space and co-visualised with the confocal stack of the clone in VVD viewer. In many cases this allowed exclusion of types definitely absent from the clone. For each clone we provide a confidence score (1: almost certain, 2: good guess, 3: putative). At the end of this process, we expanded annotation to the right hemisphere assigning clone identity to neurons from the same “group” in MANC metadata. A summary of fruitless neurons annotation is provided in Supplemental File 2.

#### Identification of sex-specific or dimorphic neurons

MANC neuronal types were considered male-specific or dimorphic if none of the neurons within the type had a FANC match with connectivity score greater than 0.3. We decided to perform a type-level analysis because it is more robust than per-neuron analysis. Due to the incomplete proofreading status of FANC it is not uncommon to obtain matches on one side but not the other. In addition, the matching algorithm can only assign a single match, which would result in unmatched neurons if a type has a different number of neurons between males and females. This level of dimorphism would need to be treated differently.

For female specific neurons in 01A T1, we identified all FANC neurons manually annotated by us to belong to that hemilineage that were not matched to any MANC neuron. We then manually curated the remainders to retain only those neurons with a clear contralateral match.

### Fly husbandry

All flies were raised in vials of iberian medium at 25°C under a 12h - 12h day-night light cycle.

## Data Availability

10x scRNA-seq data (FASTQ files, cellranger outputs and processed and annotated Seurat objects) are available at Gene Expression Omnibus (GEO) under accession number GSE304221, and at BioProject under accession number PRJNA1297747. Annotated loom files are also accessible in Zenodo (full dataset: 10.5281/zenodo.17183028; neurons: 10.5281/zenodo.17184089, glia: 10.5281/zenodo.17185431) and in SCope (https://scope.aertslab.org/#/HundredDrills/*/welcome), an interactive web platform for exploring single-cell datasets. SCope interface allows rapid inspection of marker genes and cell-level information, making it straightforward to interact with the data. Code with instructions to download and inspect data from GEO and SCope is reported in GitHub https://github.com/FlyNeuroAtlas/DevSeqVNC. Additional code and imaging data is available upon request.

## References

1. Takemura, S.-Y. et al. A Connectome of the Male Drosophila Ventral Nerve Cord. eLife 13, (2024).

2. Marin, E. C. et al. Systematic annotation of a complete adult male Drosophila nerve cord connectome reveals principles of functional organisation. (2024) doi:10.7554/elife.97766.1.

3. Sagner, A. & Briscoe, J. Establishing neuronal diversity in the spinal cord: a time and a place. Development 146, dev182154 (2019).

4. Doe, C. Q. Temporal Patterning in the Drosophila CNS. Annu Rev Cell Dev Biol 33, 219–240 (2017).

5. Jain, S. & Zipursky, S. L. Temporal control of neuronal wiring. Semin. Cell Dev. Biol. 142, 81–90 (2023).

6. Piwecka, M., Rajewsky, N. & Rybak-Wolf, A. Single-cell and spatial transcriptomics: deciphering brain complexity in health and disease. Nature Reviews Neurology 19, 346–362 (2023).

7. Zhang, M. et al. Molecularly defined and spatially resolved cell atlas of the whole mouse brain. Nature 624, 343–354 (2023).

8. Yao, Z. et al. A high-resolution transcriptomic and spatial atlas of cell types in the whole mouse brain. Nature 624, 317–332 (2023).

9. Langlieb, J. et al. The molecular cytoarchitecture of the adult mouse brain. Nature 624, 333–342 (2023).

10. Siletti, K. et al. Transcriptomic diversity of cell types across the adult human brain. Science 382, eadd7046 (2023).

11. Taylor, S. R. et al. Molecular topography of an entire nervous system. Cell 184, 4329–4347.e23 (2021).

12. Dorkenwald, S. et al. Neuronal wiring diagram of an adult brain. Nature 634, 124–138 (2024).

13. The MICrONS Consortium. Functional connectomics spanning multiple areas of mouse visual cortex. Nature 640, 435–447 (2025).

14. Wellcome Trust. Scaling up Connectomics. https://wellcome.org/reports/scaling-connectomics (2023).

15. Azevedo, A. et al. Connectomic reconstruction of a female Drosophila ventral nerve cord. Nature 631, 360–368 (2024).

16. Zeng, H. & Sanes, J. R. Neuronal cell-type classification: challenges, opportunities and the path forward. Nat. Rev. Neurosci. 18, 530–546 (2017).

17. Bates, A. S., Janssens, J., Jefferis, G. S. & Aerts, S. Neuronal cell types in the fly: single-cell anatomy meets single-cell genomics. Curr. Opin. Neurobiol. 56, 125–134 (2019).

18. Zeng, H. What is a cell type and how to define it? Cell 185, 2739–2755 (2022).

19. Schlegel, P. et al. Whole-brain annotation and multi-connectome cell typing of Drosophila. Nature 634, 139–152 (2024).

20. Stürner, T. et al. Comparative connectomics of Drosophila descending and ascending neurons. Nature (2025) doi:10.1038/s41586-025-08925-z.

21. Li, H. et al. Classifying Drosophila Olfactory Projection Neuron Subtypes by Single-Cell RNA Sequencing. Cell 171, 1206–1220.e22 (2017).

22. Simon, F., et al. High-throughput identification of the spatial origins of Drosophila optic lobe neurons using single-cell mRNA-sequencing. *bioRxivorg* (2024) doi:10.1101/2024.02.05.578975.

23. Coyne, R., et al. Regulatory logic of neuronal identity specification in Drosophila. *bioRxivorg* (2025) doi:10.1101/2025.09.01.673531.

24. Kurmangaliyev, Y. Z., Yoo, J., Valdes-Aleman, J., Sanfilippo, P. & Zipursky, S. L. Transcriptional Programs of Circuit Assembly in the Drosophila Visual System. Neuron 108, 1045–1057.e6 (2020).

25. Özel, M. N. et al. Neuronal diversity and convergence in a visual system developmental atlas. Nature 589, 88–95 (2021).

26. Özel, M. N. et al. Coordinated control of neuronal differentiation and wiring by sustained transcription factors. Science 378, eadd1884 (2022).

27. Berg, S. et al. Sexual dimorphism in the complete connectome of the Drosophila male central nervous system. bioRxivorg (2025) doi:10.1101/2025.10.09.680999.

28. Dalmaijer, E. S., Nord, C. L. & Astle, D. E. Statistical power for cluster analysis. BMC Bioinformatics 23, 205 (2022).

29. Croset, V., Treiber, C. D. & Waddell, S. Cellular diversity in the midbrain revealed by single-cell transcriptomics. Elife 7, (2018).

30. Davie, K. et al. A Single-Cell Transcriptome Atlas of the Aging Drosophila Brain. Cell 174, 982–998.e20 (2018).

31. Allen, A. M. et al. A single-cell transcriptomic atlas of the adult Drosophila ventral nerve cord. Elife 9, (2020).

32. Xie, Q. et al. Temporal evolution of single-cell transcriptomes of Drosophila olfactory projection neurons. Elife 10, (2021).

33. Truman, J. W. & Riddiford, L. M. Drosophila postembryonic nervous system development: a model for the endocrine control of development. Genetics 223, (2023).

34. Hartenstein, V. Morphological diversity and development of glia in Drosophila. Glia 59, 1237–1252 (2011).

35. Spindler, S. R. & Hartenstein, V. The Drosophila neural lineages: a model system to study brain development and circuitry. Dev. Genes Evol. 220, 1–10 (2010).

36. Etheredge, J. Transcriptional profiling of Drosophila larval ventral nervous system hemilineages. Preprint at 10.17863/CAM.17445 (2017).

37. Liu, Z. et al. Opposing intrinsic temporal gradients guide neural stem cell production of varied neuronal fates. Science 350, 317–320 (2015).

38. Nguyen, T. H. et al. scRNA-seq data from the larval Drosophila ventral cord provides a resource for studying motor systems function and development. Dev Cell 59, 1210–1230.e9 (2024).

39. Lacin, H. & Truman, J. W. Lineage mapping identifies molecular and architectural similarities between the larval and adult Drosophila central nervous system. Elife 5, e13399 (2016).

40. Truman, J. W., Moats, W., Altman, J., Marin, E. C. & Williams, D. W. Role of Notch signaling in establishing the hemilineages of secondary neurons in Drosophila melanogaster. Development 137, 53–61 (2010).

41. Lacin, H. et al. Neurotransmitter identity is acquired in a lineage-restricted manner in the CNS. Elife 8, (2019).

42. Eckstein, N., et al. Neurotransmitter Classification from Electron Microscopy Images at Synaptic Sites in Drosophila Melanogaster. Cell in press, (2024).

43. Soffers, J. H. M. et al. A library of lineage-specific driver lines connects developing neuronal circuits to behavior in the Drosophila ventral nerve cord. Elife 14, (2025).

44. Truman, J. W., Schuppe, H., Shepherd, D. & Williams, D. W. Developmental architecture of adult-specific lineages in the ventral CNS of Drosophila. Development 131, 5167–5184 (2004).

45. Marin, E. C. et al. Ultrabithorax confers spatial identity in a context-specific manner in the Drosophila postembryonic ventral nervous system. Neural Dev. 7, 31 (2012).

46. Kuert, P. A., Hartenstein, V., Bello, B. C., Lovick, J. K. & Reichert, H. Neuroblast lineage identification and lineage-specific Hox gene action during postembryonic development of the subesophageal ganglion in the Drosophila central brain. Dev. Biol. 390, 102–115 (2014).

47. Shepherd, D., Sahota, V., Court, R., Williams, D. W. & Truman, J. W. Developmental organization of central neurons in the adult Drosophila ventral nervous system. J. Comp. Neurol. 527, 2573–2598 (2019).

48. Lin, S., Kao, C.-F., Yu, H.-H., Huang, Y. & Lee, T. Lineage analysis of Drosophila lateral antennal lobe neurons reveals notch-dependent binary temporal fate decisions. PLoS Biol. 10, e1001425 (2012).

49. Lee, Y.-J. et al. Conservation and divergence of related neuronal lineages in the Drosophila central brain. Elife 9, (2020).

50. Li, X. et al. Temporal patterning of Drosophila medulla neuroblasts controls neural fates. Nature 498, 456–462 (2013).

51. Marquardt, T. & Gruss, P. Generating neuronal diversity in the retina: one for nearly all. Trends Neurosci 25, 32–38 (2002).

52. Guan, W. et al. Post-transcriptional regulation of transcription factor codes in immature neurons drives neuronal diversity. Cell Rep. 39, 110992 (2022).

53. Zhou, B., Williams, D. W., Altman, J., Riddiford, L. M. & Truman, J. W. Temporal patterns of broad isoform expression during the development of neuronal lineages in Drosophila. Neural Dev. 4, 39 (2009).

54. Trapnell, C. et al. The dynamics and regulators of cell fate decisions are revealed by pseudotemporal ordering of single cells. Nat. Biotechnol. 32, 381–386 (2014).

55. Qiu, X. et al. Reversed graph embedding resolves complex single-cell trajectories. Nat. Methods 14, 979–982 (2017).

56. Cao, J. et al. The single-cell transcriptional landscape of mammalian organogenesis. Nature 566, 496–502 (2019).

57. Traag, V. A., Waltman, L. & van Eck, N. J. From Louvain to Leiden: guaranteeing well-connected communities. Sci. Rep. 9, 5233 (2019).

58. Levine, J. H. et al. Data-driven phenotypic dissection of AML reveals progenitor-like cells that correlate with prognosis. Cell 162, 184–197 (2015).

59. Sumanaweera, D. et al. Gene-level alignment of single-cell trajectories. Nat. Methods (2024) doi:10.1038/s41592-024-02378-4.

60. Pollington, H. Q., Seroka, A. Q. & Doe, C. Q. From temporal patterning to neuronal connectivity in Drosophila type I neuroblast lineages. Semin. Cell Dev. Biol. 142, 4–12 (2023).

61. Isshiki, T., Pearson, B., Holbrook, S. & Doe, C. Q. Drosophila neuroblasts sequentially express transcription factors which specify the temporal identity of their neuronal progeny. Cell 106, 511–521 (2001).

62. Hobert, O. Terminal selectors of neuronal identity. Curr. Top. Dev. Biol. 116, 455–475 (2016).

63. Urbach, R., Jussen, D. & Technau, G. M. Gene expression profiles uncover individual identities of gnathal neuroblasts and serial homologies in the embryonic CNS of Drosophila. Development 143, 1290–1301 (2016).

64. Lillvis, J. L. et al. Nested neural circuits generate distinct acoustic signals during Drosophila courtship. Curr. Biol. 34, 808–824.e6 (2024).

65. Shirangi, T. R., Wong, A. M., Truman, J. W. & Stern, D. L. Doublesex regulates the connectivity of a neural circuit controlling Drosophila male courtship song. Dev. Cell 37, 533–544 (2016).

66. Rezával, C., Nojima, T., Neville, M. C., Lin, A. C. & Goodwin, S. F. Sexually dimorphic octopaminergic neurons modulate female postmating behaviors in Drosophila. Curr. Biol. 24, 725–730 (2014).

67. Dann, E., Henderson, N. C., Teichmann, S. A., Morgan, M. D. & Marioni, J. C. Differential abundance testing on single-cell data using k-nearest neighbor graphs. Nat. Biotechnol. 40, 245–253 (2022).

68. Asahina, K. Sex differences in Drosophila behavior: Qualitative and Quantitative Dimorphism. Curr. Opin. Physiol. 6, 35–45 (2018).

69. Cachero, S., Ostrovsky, A. D., Yu, J. Y., Dickson, B. J. & Jefferis, G. S. X. E. Sexual dimorphism in the fly brain. Curr. Biol. 20, 1589–1601 (2010).

70. Kimura, K.-I., Ote, M., Tazawa, T. & Yamamoto, D. Fruitless specifies sexually dimorphic neural circuitry in the Drosophila brain. Nature 438, 229–233 (2005).

71. Sanders, L. E. & Arbeitman, M. N. Doublesex establishes sexual dimorphism in the Drosophila central nervous system in an isoform-dependent manner by directing cell number. Dev. Biol. 320, 378–390 (2008).

72. Ren, Q., Awasaki, T., Huang, Y.-F., Liu, Z. & Lee, T. Cell Class-Lineage Analysis Reveals Sexually Dimorphic Lineage Compositions in the Drosophila Brain. Curr Biol 26, 2583–2593 (2016).

73. Sanes, J. R. & Zipursky, S. L. Synaptic specificity, recognition molecules, and assembly of neural circuits. Cell 181, 536–556 (2020).

74. Elkahlah, N., Lin, Y., Shirangi, T. R. & Clowney, E. J. Hierarchical diversification of instinctual behavior neurons by lineage, birth order, and sex. bioRxivorg (2025) doi:10.1101/2025.06.03.657692.

75. Allen, A. M., Neville, M. C., Nojima, T., Alejevski, F. & Goodwin, S. F. Differential neuronal survival defines a novel axis of sexual dimorphism in the Drosophila brain. Cell Genom. 101125 (2026) doi:10.1016/j.xgen.2025.101125.

76. Li, Z., et al. Repulsive interactions instruct synaptic partner matching in an olfactory circuit. *bioRxivorg* (2025) doi:10.1101/2025.03.01.640985.

77. Yoo, J. et al. Brain wiring determinants uncovered by integrating connectomes and transcriptomes. Curr. Biol. 33, 3998–4005.e6 (2023).

78. Carrier, Y. et al. Biased cell adhesion organizes the Drosophila visual motion integration circuit. Dev. Cell 60, 762–779.e7 (2025).

79. Sanfilippo, P. et al. Mapping of multiple neurotransmitter receptor subtypes and distinct protein complexes to the connectome. Neuron 112, 942–958.e13 (2024).

80. Yu, H.-H. et al. A complete developmental sequence of a Drosophila neuronal lineage as revealed by twin-spot MARCM. PLoS Biol. 8, e1000461 (2010).

81. Tavakoli, M. R. et al. Light-microscopy-based connectomic reconstruction of mammalian brain tissue. Nature 642, 398–410 (2025).

82. Stanley, G., Gokce, O., Malenka, R. C., Südhof, T. C. & Quake, S. R. Continuous and discrete neuron types of the adult Murine striatum. Neuron 105, 688–699.e8 (2020).

83. Ren, Q. et al. Stem cell-intrinsic, Seven-up-triggered temporal factor gradients diversify intermediate neural progenitors. Curr. Biol. 27, 1303–1313 (2017).

84. Konstantinides, N. et al. A complete temporal transcription factor series in the fly visual system. Nature 604, 316–322 (2022).

85. Lee, J. Y., Huang, N., Samuels, T. J. & Davis, I. Imp/IGF2BP and Syp/SYNCRIP temporal RNA interactomes uncover combinatorial networks of regulators of brain development. Sci Adv 11, eadr6682 (2025).

86. Cruz, J., Ureña, E., Franch-Marro, X. & Martín, D. E93 controls adult differentiation by repressingbroadinDrosophila. *bioRxiv* (2024) doi:10.1101/2024.02.14.580315.

87. Smolin, N. et al. Neuronal identity control at the resolution of a single transcription factor isoform. bioRxivorg (2024) doi:10.1101/2024.06.14.598883.

88. Farnworth, M. S., Bucher, G. & Hartenstein, V. An atlas of the developing Tribolium castaneum brain reveals conservation in anatomy and divergence in timing to Drosophila melanogaster. J. Comp. Neurol. (2022) doi:10.1002/cne.25335.

89. Hasegawa, E. et al. Concentric zones, cell migration and neuronal circuits in the Drosophila visual center. Development 138, 983–993 (2011).

90. Ngo, K. T., Andrade, I. & Hartenstein, V. Spatio-temporal pattern of neuronal differentiation in the Drosophila visual system: A user’s guide to the dynamic morphology of the developing optic lobe. Dev. Biol. 428, 1–24 (2017).

91. Allen, A. M. et al. A high-resolution atlas of the brain predicts lineage and birth order underly neuronal identity. bioRxiv (2025) doi:10.1101/2025.06.04.657818.

92. Mori, H. et al. Expression of mouse igf2 mRNA-binding protein 3 and its implications for the developing central nervous system. J Neurosci Res 64, 132–143 (2001).

93. Chen, H.-H. et al. hnRNP Q regulates Cdc42-mediated neuronal morphogenesis. Mol Cell Biol 32, 2224–2238 (2012).

94. Sagner, A. et al. A shared transcriptional code orchestrates temporal patterning of the central nervous system. PLoS Biol. 19, e3001450 (2021).

95. Osseward, P. J., 2nd et al. Conserved genetic signatures parcellate cardinal spinal neuron classes into local and projection subsets. Science 372, 385–393 (2021).

96. Kishore, S., Cadoff, E. B., Agha, M. A. & McLean, D. L. Orderly compartmental mapping of premotor inhibition in the developing zebrafish spinal cord. Science 370, 431–436 (2020).

97. Huszár, R., Zhang, Y., Blockus, H. & Buzsáki, G. Preconfigured dynamics in the hippocampus are guided by embryonic birthdate and rate of neurogenesis. Nat Neurosci 25, 1201–1212 (2022).

98. Dong, P. D. S., Dicks, J. S. & Panganiban, G. Distal-less and homothorax regulate multiple targets to pattern the Drosophila antenna. Development 129, 1967–1974 (2002).

99. Emerald, B. S., Curtiss, J., Mlodzik, M. & Cohen, S. M. Distal antenna and distal antenna related encode nuclear proteins containing pipsqueak motifs involved in antenna development in Drosophila. Development 130, 1171–1180 (2003).

100. Baanannou, A. et al. Drosophila distal-less and Rotund bind a single enhancer ensuring reliable and robust bric-a-brac2 expression in distinct limb morphogenetic fields. PLoS Genet 9, e1003581 (2013).

101. Mackay, T. F. C. et al. The Drosophila melanogaster Genetic Reference Panel. Nature 482, 173–178 (2012).

102. Huang, Y., McCarthy, D. J. & Stegle, O. Vireo: Bayesian demultiplexing of pooled single-cell RNA-seq data without genotype reference. Genome Biol. 20, 273 (2019).

103. 103.[No title]. https://ftp.ensembl.org/pub/release-109/fasta/drosophila_melanogaster/dna/Drosophila_melanogaster.BDGP6.32.dna_rm.toplevel.fa.gz

104. Neavin, D. et al. Demuxafy: improvement in droplet assignment by integrating multiple single-cell demultiplexing and doublet detection methods. Genome Biol. 25, 94 (2024).

105. Seroka, A., Lai, S.-L. & Doe, C. Q. Transcriptional profiling from whole embryos to single neuroblast lineages in Drosophila. Dev. Biol. 489, 21–33 (2022).

106. Integrated analysis of multimodal single-cell data. Cell 184, 3573–3587.e29 (2021).

107. Hao, Y. et al. Dictionary learning for integrative, multimodal and scalable single-cell analysis. Nat Biotechnol 42, 293–304 (2024).

108. Lago-Baldaia, I. et al. A Drosophila glial cell atlas reveals a mismatch between transcriptional and morphological diversity. PLoS Biol. 21, e3002328 (2023).

109. Galdos, F. X. et al. devCellPy is a machine learning-enabled pipeline for automated annotation of complex multilayered single-cell transcriptomic data. Nat. Commun. 13, 5271 (2022).

110. Tirosh, I. et al. Dissecting the multicellular ecosystem of metastatic melanoma by single-cell RNA-seq. Science 352, 189–196 (2016).

111. Pfeiffer, B. D. et al. Refinement of tools for targeted gene expression in Drosophila. Genetics 186, 735–755 (2010).

112. Awasaki, T. et al. Making Drosophila lineage-restricted drivers via patterned recombination in neuroblasts. Nat. Neurosci. 17, 631–637 (2014).

113. Marin, E. C. et al. Systematic annotation of a complete adult male Drosophila nerve cord connectome reveals principles of functional organisation. bioRxiv 2023.06.05.543407 (2023) doi:10.1101/2023.06.05.543407.

114. Soffers, J. H. M. et al. A library of lineage-specific driver lines connects developing neuronal circuits to behavior in the ventral nerve cord. Elife 14, (2025).

115. Li, H.-H. et al. A GAL4 driver resource for developmental and behavioral studies on the larval CNS of Drosophila. Cell Rep. 8, 897–908 (2014).

116. Qiu, X. et al. Single-cell mRNA quantification and differential analysis with Census. Nat. Methods 14, 309–315 (2017).

117. Ostrovsky, A., Cachero, S. & Jefferis, G. Clonal analysis of olfaction in Drosophila: immunochemistry and imaging of fly brains. Cold Spring Harb. Protoc. 2013, 342–346 (2013).

118. Lee, T. & Luo, L. Mosaic analysis with a repressible cell marker for studies of gene function in neuronal morphogenesis. Neuron 22, 451–461 (1999).

119. Macabenta, F. & Stathopoulos, A. Migrating cells control morphogenesis of substratum serving as track to promote directional movement of the collective. Development 146, dev177295 (2019).

120. Jinushi-Nakao, S. et al. Knot/Collier and cut control different aspects of dendrite cytoskeleton and synergize to define final arbor shape. Neuron 56, 963–978 (2007).

121. Daul, A. L., Komori, H. & Lee, C.-Y. EdU (5-ethynyl-2’-deoxyuridine) labeling of Drosophila mitotic neuroblasts. Cold Spring Harb. Protoc. 2010, db.prot5461 (2010).

122. Preibisch, S., Saalfeld, S. & Tomancak, P. Globally optimal stitching of tiled 3D microscopic image acquisitions. Bioinformatics 25, 1463–1465 (2009).

123. Berg, S. et al. Ilastik: Interactive machine learning for (bio)image analysis. Nat. Methods 16, 1226–1232 (2019).

124. Bates, A. S. et al. The natverse, a versatile toolbox for combining and analysing neuroanatomical data. Elife 9, (2020).

125. Costa, M., Manton, J. D., Ostrovsky, A. D., Prohaska, S. & Jefferis, G. S. X. E. NBLAST: Rapid, Sensitive Comparison of Neuronal Structure and Construction of Neuron Family Databases. Neuron 91, 293–311 (2016).

126. GitHub - flyconnectome/fancr: Support Acccess to the Female Adult Nerve Cord (FANC) Dataset. *GitHub* https://github.com/flyconnectome/fancr.

