## Extended Data Figures 1-13 for "A Developmental Atlas of the *Drosophila* Nerve Cord Uncovers a Global Temporal Code for Neuronal Identity"

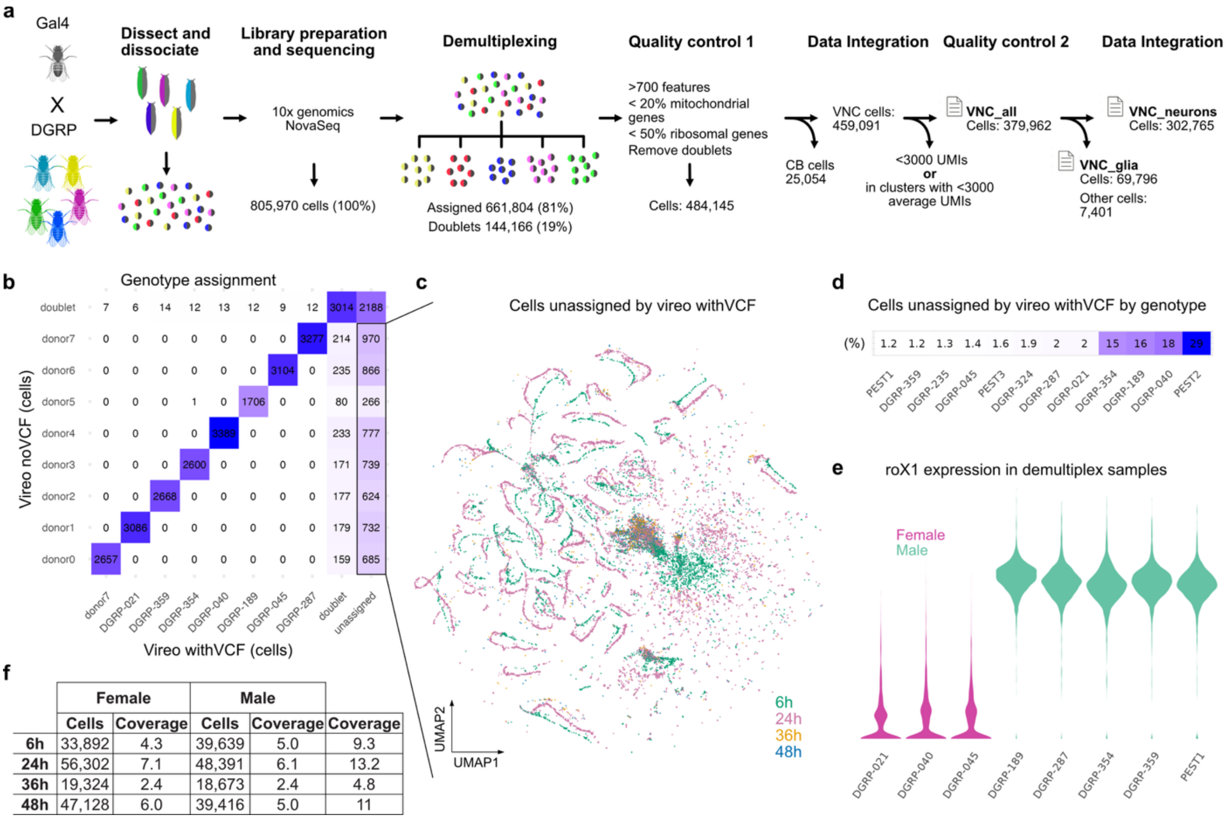


| | Female | | Male | | |
|---|---|---|---|---|---|
| | Cells | Coverage | Cells | Coverage | Coverage |
| 6h | 33,892 | 4.3 | 39,639 | 5.0 | 9.3 |
| 24h | 56,302 | 7.1 | 48,391 | 6.1 | 13.2 |
| 36h | 19,324 | 2.4 | 18,673 | 2.4 | 4.8 |
| 48h | 47,128 | 6.0 | 39,416 | 5.0 | 11 |

**Extended Data Fig. 1 | Generation of the scRNAseq dataset**

**a**, Data acquisition and processing pipeline. Flies from Gal4 and DGRP lines were crossed, staged, dissected and dissociated. Cell suspensions were used to make 10X genomics libraries and sequenced. Cellranger analysis generated 805,970 cells, resulting in 661,804 singlets after demultiplexing using vireo. 484,145 cells passed the first quality control step. After removing 25,054 cells from unrelated brain samples we merged samples from different genotypes using Seurat and removed non-neuronal cells. Following a second quality control step, 302,765 neurons were left, which form the final dataset used. **b**, Confusion matrix for one sample comparing Vireo demultiplexing results with and without a Variant Call Format (VCF) file. Box marks cells assigned to a group only in the run without the VCF, included in (c). **c**, UMAP plot showing the distribution of unassigned neurons by Vireo run with VCF, but assigned by Vireo without VCF reference across all experiments. **d,** Percentage of cells per genotype that were unassigned by vireo, when used with a VCF, but assigned when not using a VCF reference, showing a genotype bias. **e**, Violin plots showing expression level of the long non-coding RNA roX1, a marker for male cells, in the cells from (b). Violin colour indicates the sex of the corresponding dissected animals. Correspondence between high roX1 level and male animal identity indicates the quality of the demultiplexing results. **f,** Table showing the number of neurons passing QC by stage and sex, along with coverage estimates based on the 7,880 non-glial, soma-positioned neurons per side in the male connectome (MANC).

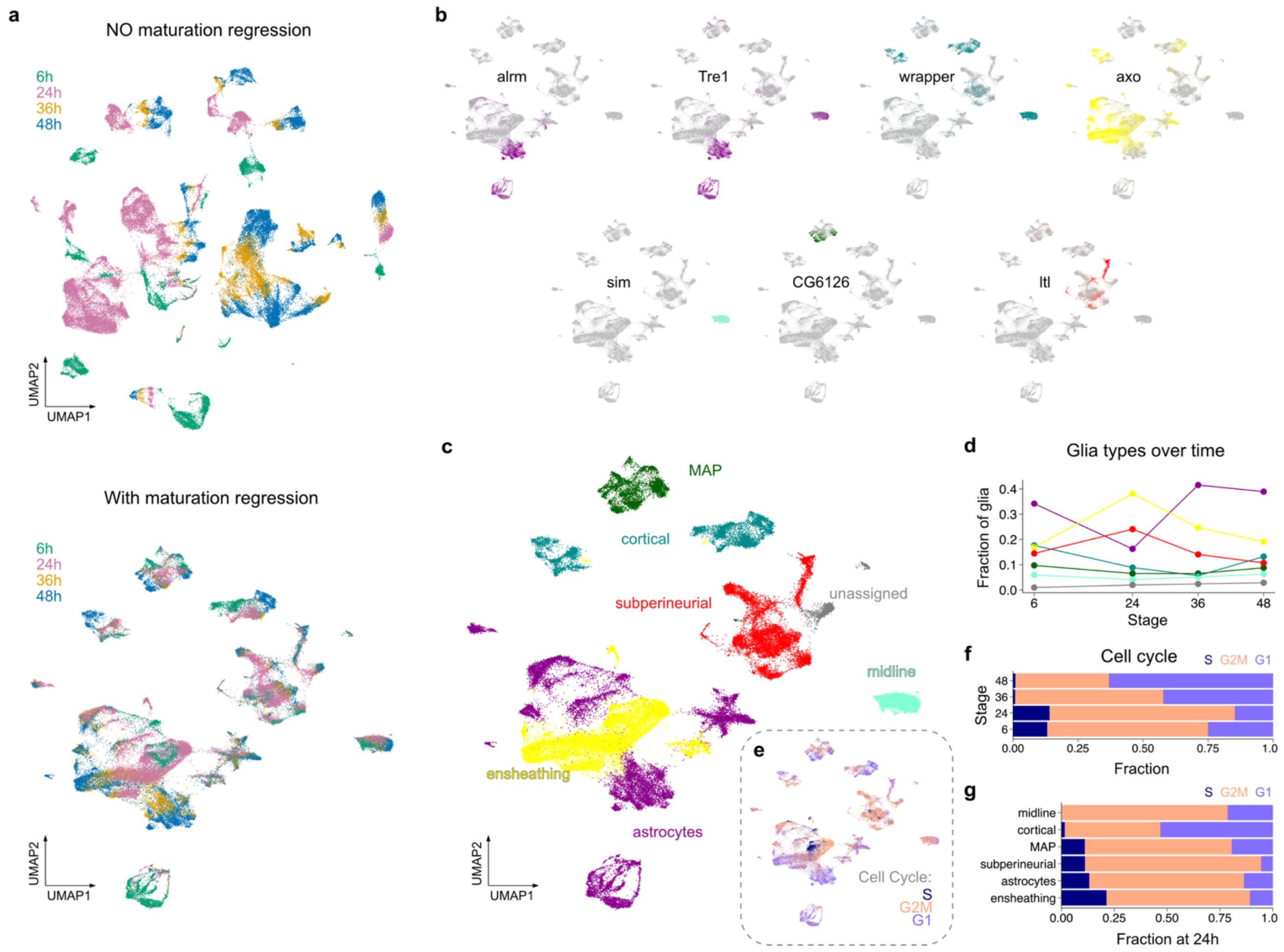


**Extended Data Fig. 2 | Developmental atlas of VNC glia**

**a,** UMAP plot of VNC glia from 6, 24, 36 and 48h without (top) and with (bottom, used in the rest of the plots) regression of maturation-driven variability. Cells are coloured by the developmental stage of the animal they originate from. **b,** UMAP plots showing expression pattern of known markers of glia cell types. **c,** UMAP plot showing glia cell types annotation based on markers in (b). **d,** Relative abundance of glia types over time. Colours as in (c). **e,** UMAP plot showing the result of cell cycle phase analysis. **f-g**, Stacked bar chart showing the fraction of glial cells in each cell cycle phase over time (f) and per glia type at 24h (g).

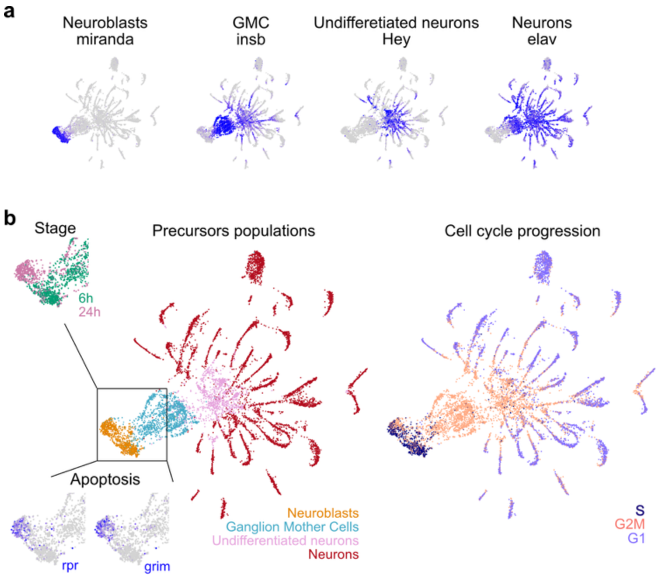


**Extended Data Fig. 3 | Annotation of neuronal progenitors**

**a,** UMAP embedding of precursor cells from Fig. 1c showing the expression of marker genes for neuroblasts, ganglion mother cells (GMC), undifferentiated neurons and differentiating neurons. **b,** UMAP plot of precursors coloured by assigned precursor category (left) and predicted cell cycle phase (right). Insets show expression of *rpr* and *grim* in neuroblasts (bottom) according to stage.

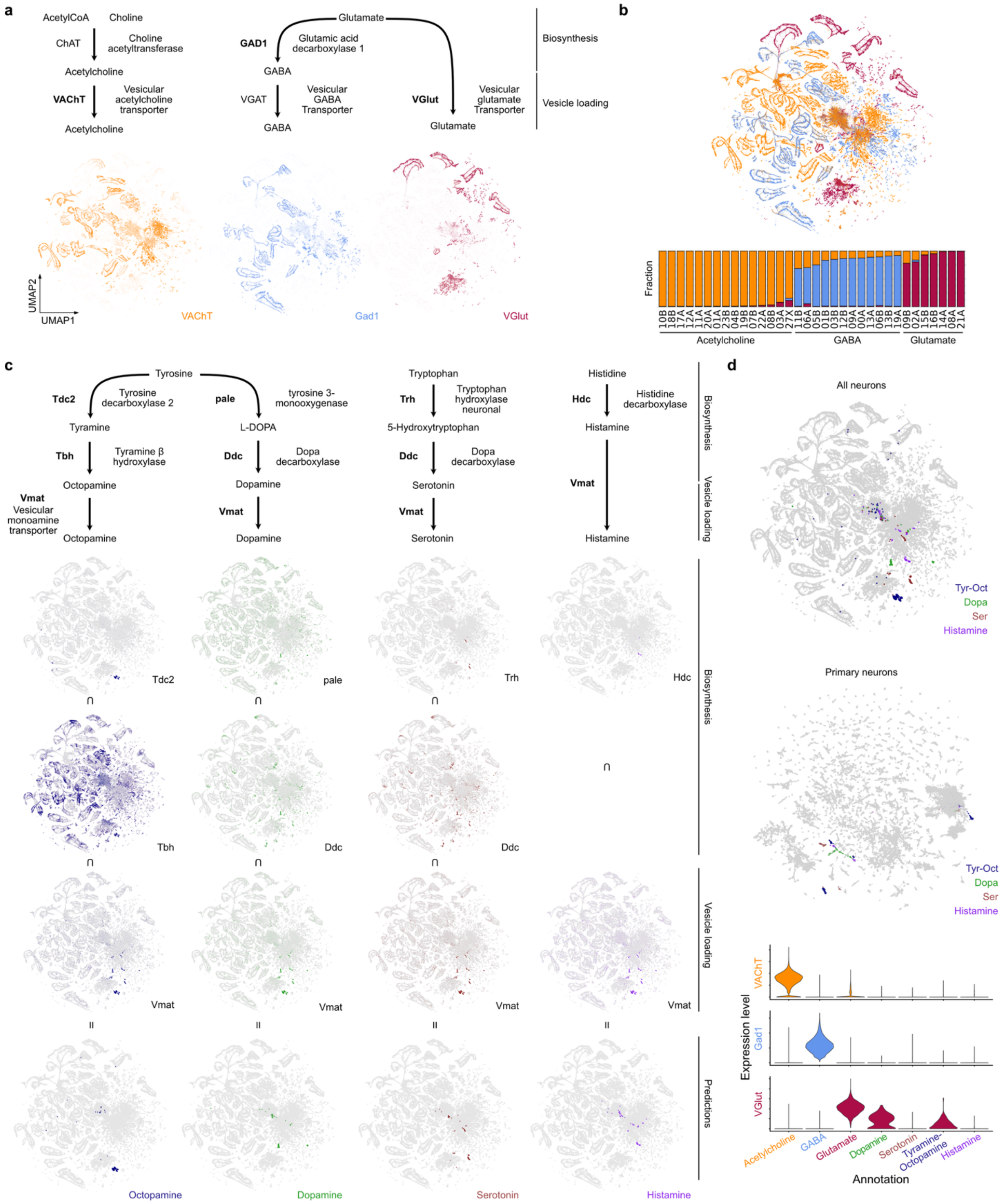


**Extended Data Fig. 4 | Predictions for neurotransmitter usage**

**a**, Top: Fast neurotransmitters biosynthetic pathways. Bottom: UMAP showing expression levels of enzymes used to predict neurotransmitter usage. **b**, Top: UMAP showing fast neurotransmitter assignments defined by the most highly expressed enzyme in (a) after scaling. Bottom: fraction of cells using each neurotransmitter system per hemillineage. **c**, Top: Aminergic neurotransmitter biosynthetic pathways. Bottom: UMAP showing expression level of enzymes and the shared vesicular monoamine transporter (Vmat) used for synaptic loading. **d,** Top: UMAP plots showing aminergic predictions resulting from intersection of biosynthetic enzymes and transporter expressions for all neurons and primary neurons only. Bottom: Violin plots show the expression of fast neurotransmitter enzymes and transporters in neurons using different fast and aminergic neurotransmitters.

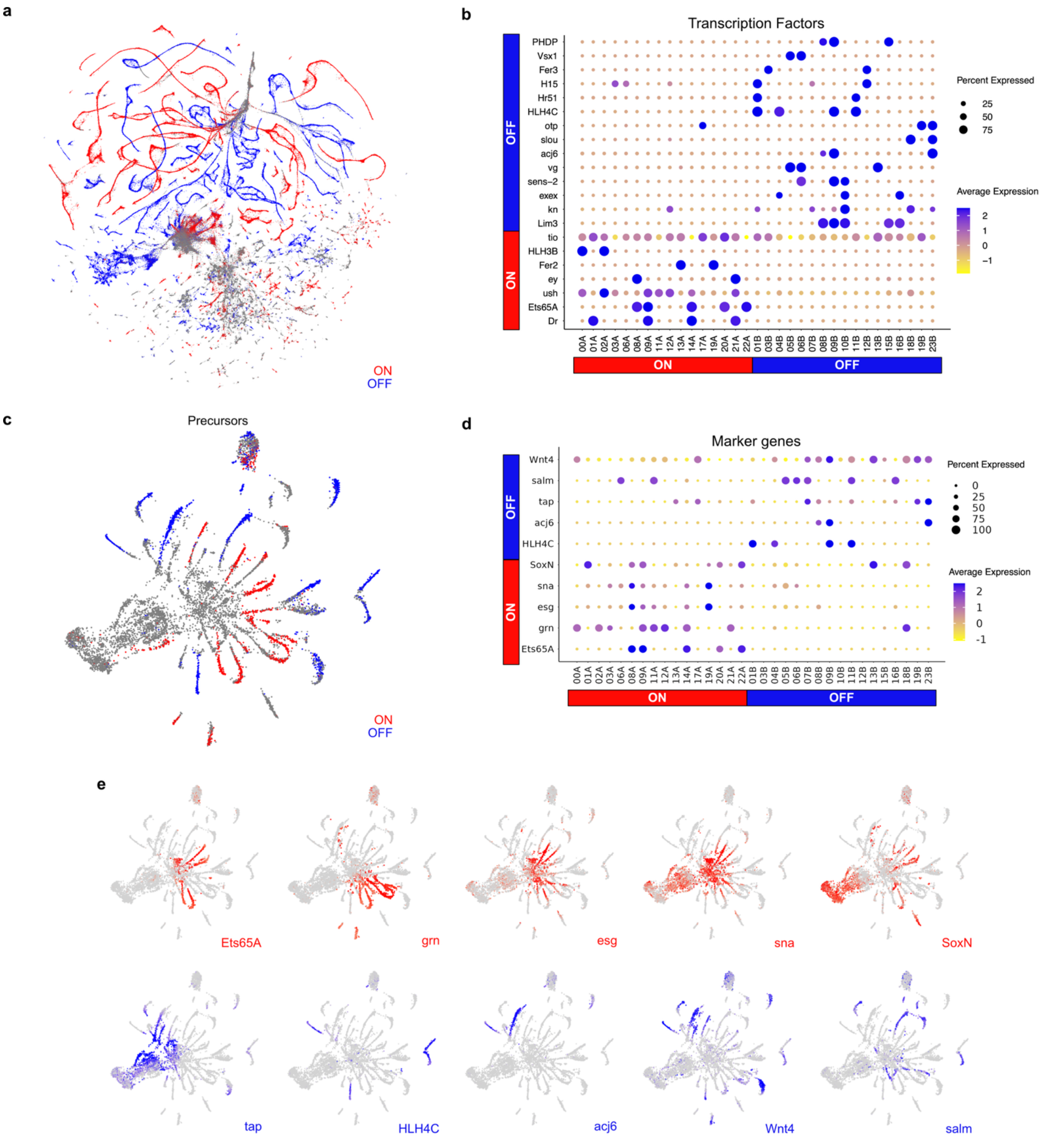


**Extended Data Fig. 5 | Markers correlated to Notch signalling**

**a,** UMAP plot showing the identity of Notch ON (A) and Notch OFF (B) hemilineages. **b,** Dot-plot showing expression in hemilineages of TFs identified as expressed preferentially in cells from Notch ON vs Notch OFF hemilineages. **c,** Same as (a) but in precursor cells (6 and 24h). **d,** Same as (b) but in precursors. Analysis expanded to all markers, not only TFs. **e,** UMAP plots showing expression of the top 5 markers for Notch ON and Notch OFF neurons in precursor cells.

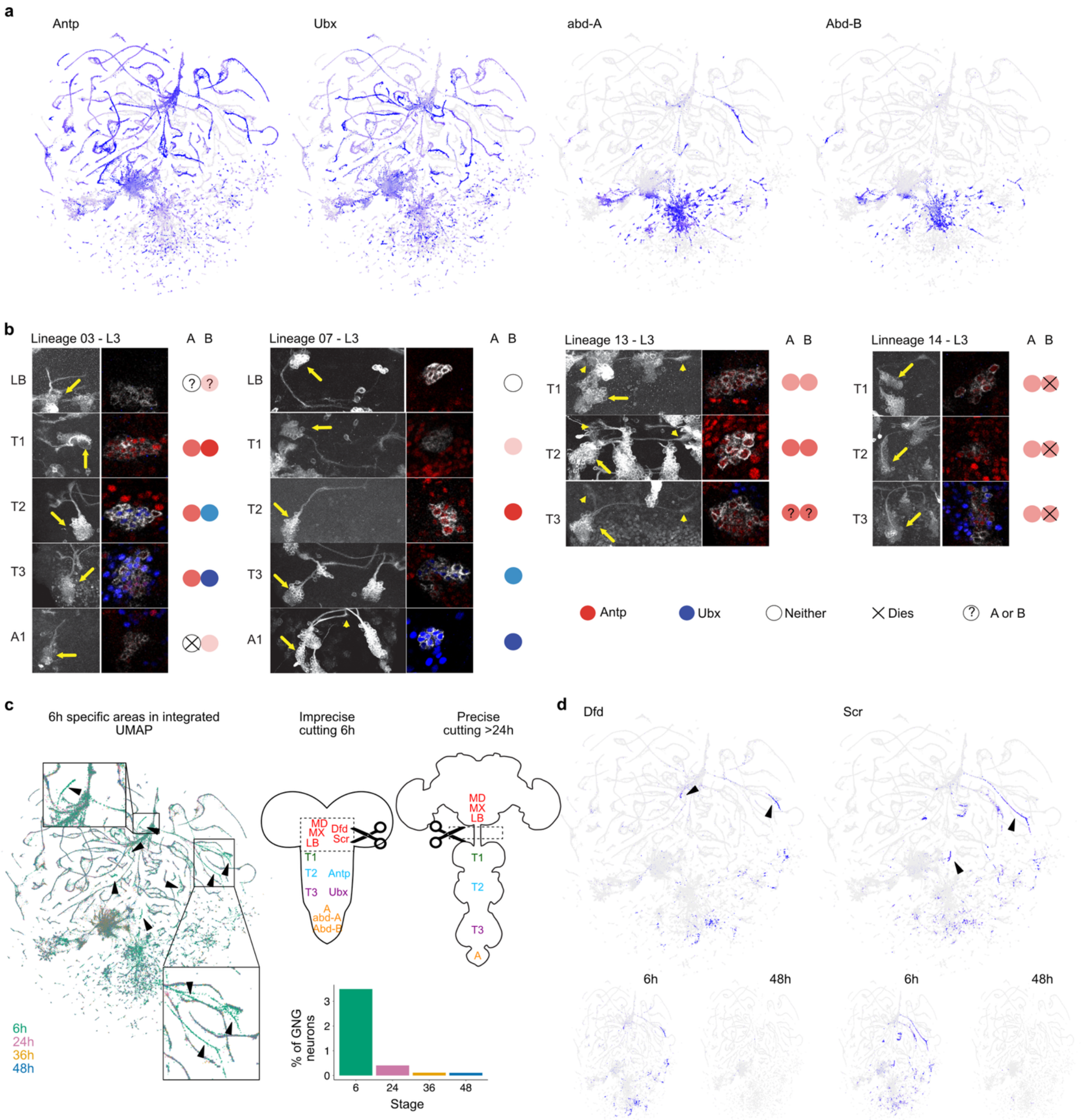


### Extended Data Fig. 6 | Expression of Hox genes

**a**, UMAP plots showing Hox genes expression across the dataset. **b,** Double staining for Antp and Ubx showing their protein levels across segments in specific lineages labelled by MARCM clones in late L3 larvae. Yellow arrows point to the somas of the lineages under consideration that can be identified by the location of the primary neurite. Images show partial max intensity projections. **c**, Left: UMAP plot coloured by to developmental stage. Insets and arrowheads highlight 6h-specific locations outside the stem region. Top right: CNS silhouette at 6 and 24-48h. The established model of the correspondence between segment and Hox gene expression is indicated. Dashed rectangles and scissors indicate areas of cutting. GNG = gnathal ganglia, LB = labial, MX = maxillary, MD = mandibular. **d,** UMAP plots showing expression of Hox genes specific for the GNG (*Dfd*, *Scr*) in all developmental stages (above) and split by stage (below) to highlight expression at 6h, but not at 48h.

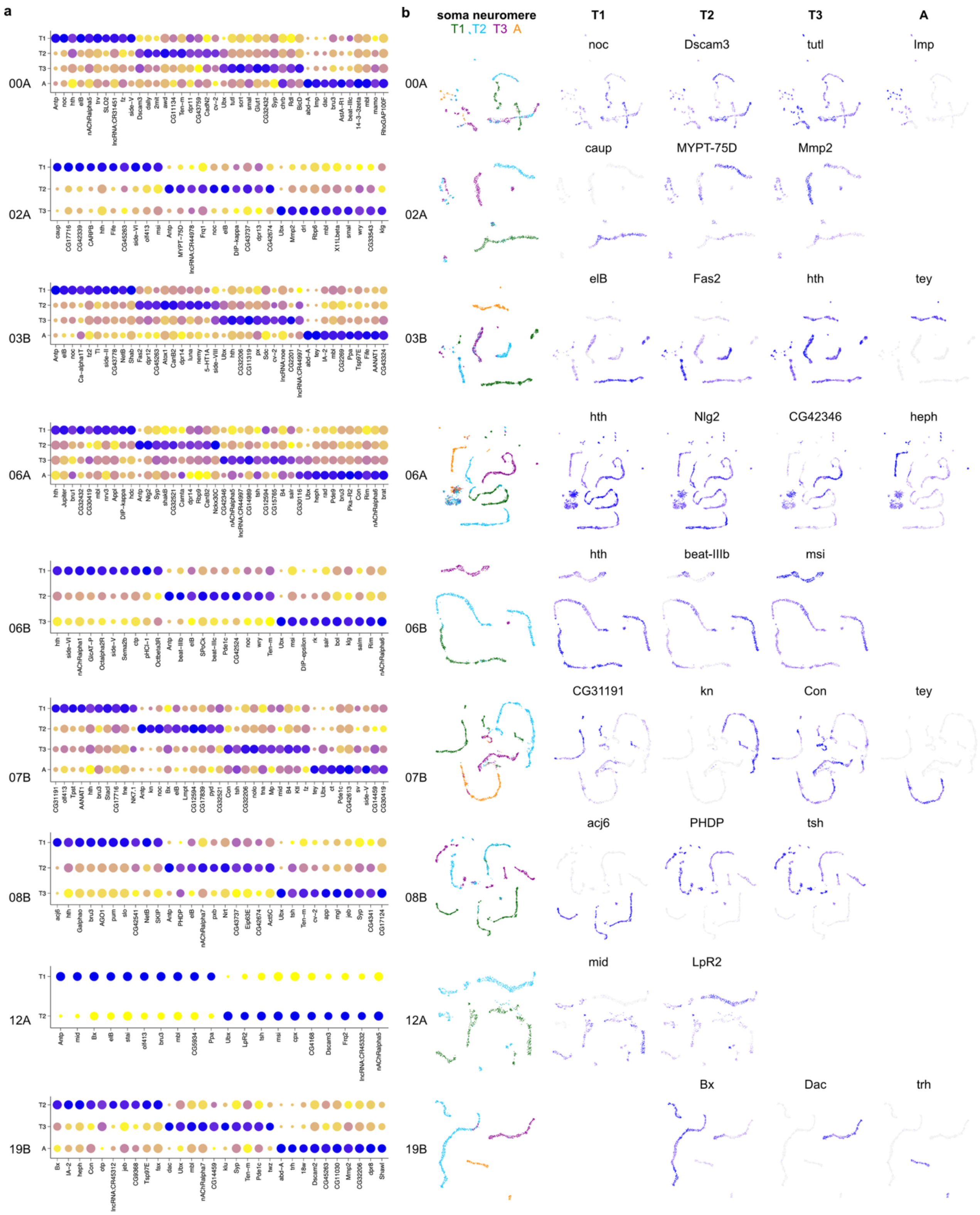


**Extended Data Fig. 7 | Segment specific markers in complex hemilineages**

**a**, Dot plots showing the expression of the top 10 segment-specific markers for hemilineages with complex UMAP trajectories. **b**, Hemilineage-specific UMAP plots showing soma segment annotation of 48h secondary neurons and the most significant segment-specific marker.

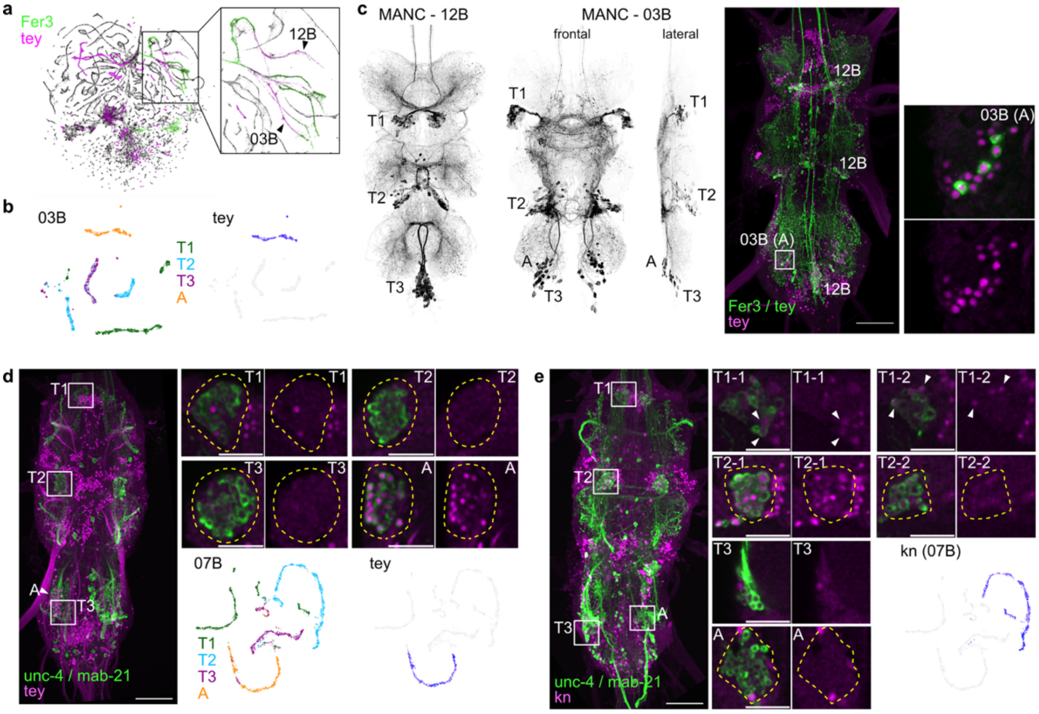


**Extended Data Fig. 8 | Validation of segments annotation**

**a,** UMAP showing co-expression of *Fer3* and *tey* in secondary hemilineages 12B and 03B. **b,** UMAPs showing 03B secondary neurons (48h only) upon re-clustering. Colour coding: segment annotation (left) and *tey* expression levels (right). Only abdominal neurons are predicted to be *tey*+. **c,** Greyscale: MANC neuronal meshes for 12B (left, frontal view) and 03B (middle, frontal and lateral views) secondary neurons. Right: MIP of confocal z-stack showing split-GAL4 intersection pattern of *Fer3* and *tey* (green) and tey protein (magenta). For 03B hemilineage, labelling is restricted to abdominal neurons which express tey. **d,** Split-GAL4 intersection of *unc-4* and *mab-21* (green), specifically labelling 07B hemilineage across segments, and tey protein (magenta). Arrowhead points to abdominal neurons that are posterior and therefore partially overlapping with T3 ones in the MIP. Insets show 07B neuronal somas across segments, with tey expression restricted to the abdominal segment as predicted by the atlas. UMAPs show 07B secondary neurons (48h only) upon re-clustering. Colour coding: segment annotation (left) and *tey* expression levels (right). **e,** Split-GAL4 intersection of *unc-4* and *mab-21* (green) and kn protein (magenta). Insets show 07B neuronal somas across segments, with kn expression restricted to a subset of T1 neurons (arrowheads) and most T2 neurons, as predicted by the atlas (see UMAP for 07B). Insets show a single z-plane; in e two planes are shown for T1 and T2 to show kn expression heterogeneity. Scale bars: MIPs = 50 µm; insets: 20 µm.

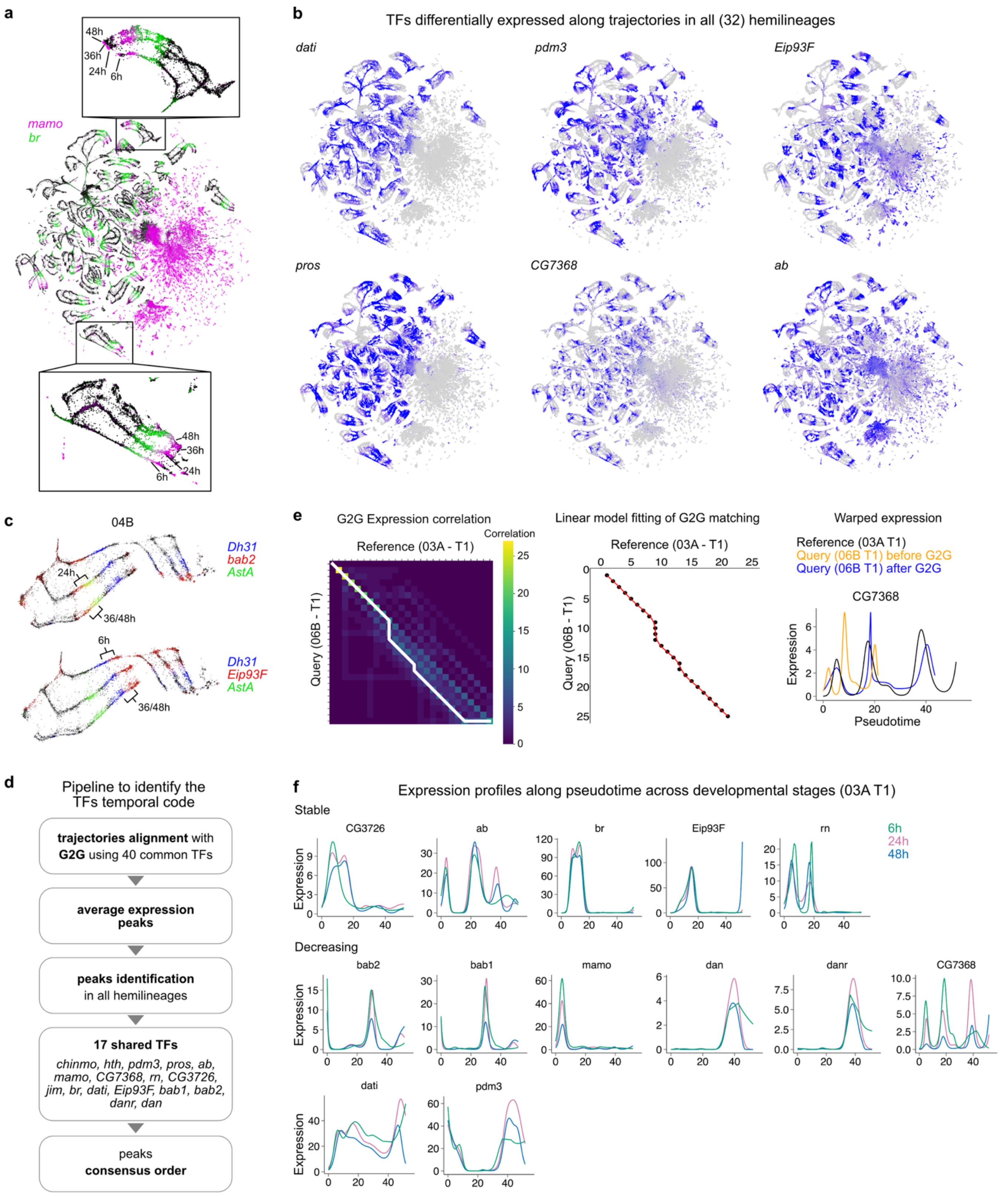


**Extended Data Fig. 9 | Shared TFs expression across development**

**a,** UMAP plots showing expression of TFs with known birth time bias, *mamo* (magenta) and *br* (green), expressed subsequently along hemilineage trajectories across developmental stages. Insets zoom on two specific hemilineages. **b,** UMAP plots showing striped expression along hemilineage trajectories of TFs differentially expressed in all hemilineages. **c,** UMAP plots of hemilineage 04B showing expression of a shTF (*bab1* above and *Eip93F* below, shown in red) and two neuropeptides (*Dh31* in blue and *AstA* in red) across developmental stages, highlighting the conserved relative positioning of the shTFs to neuropeptide-specific cells across stages. **d,** Pipeline to identify TFs with shared peaks of expression across trajectories. **e,** Left: heatmap showing bin-wide expression correlation between query (06B-T1) and reference (03A-T1) trajectories, output of G2G analysis. Middle: linear model (red line) fitting of corresponding bins from G2G alignment defining the warping function to align 06B-T1 pseudotimes to 03A-T1 reference. Right: fitted *CG7468* expression profile along pseudotime for reference (03A-T1, black), query hemilineage (06B-T1) before (yellow) and after (blue) G2G based alignment. **f,** shTFs expression along pseudotime and across stages in 03A-T1 divided according to the stability of expression between 24 and 48h. All UMAP plots are without stage variability regression.

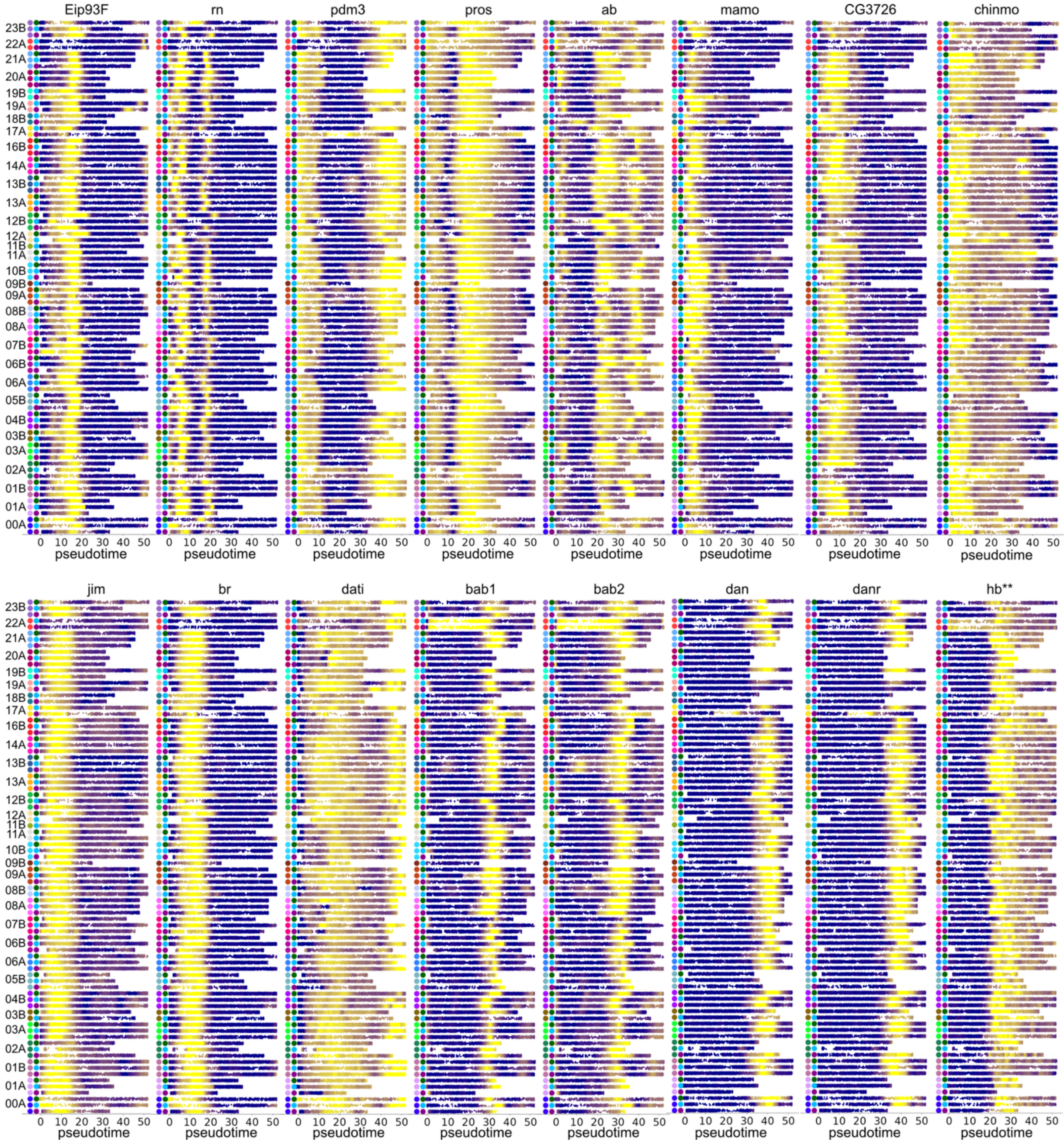


**Extended Data Fig. 10 | Global patterning of shTFs expression across hemilineages**

Normalised expression along aligned pseudotime of shTFs. Each row represents a hemilineage-segment trajectory. Colours on the left refer to hemilineage (left, refer to Fig. 2c for colour correspondence) and segment (right, refer to Fig. 3c for colour correspondence). Hb, the first temporal TF in the embryonic neuroblast temporal cascade, is also shown. Despite not emerging from our analysis of DEGs along trajectories, hb shows a consistent patterned expression across trajectories.

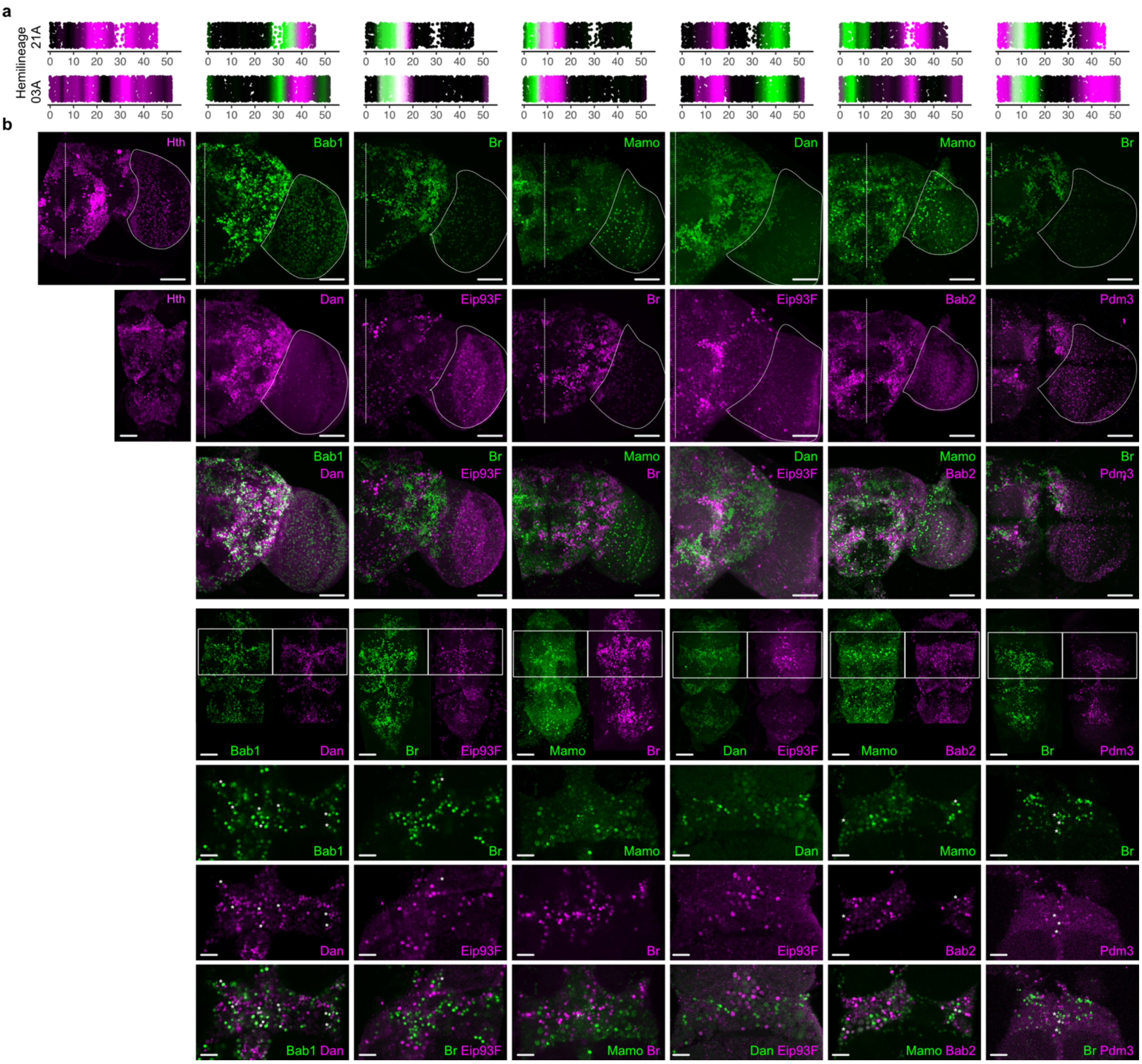


**Extended Data Fig. 11 | Validation of shared TFs expression**

**a**, Scatter plots of normalised shTFs expression in hemilineages 21A and 03A along warped pseudotime. shTF and colour correspond to columns in b. **b,** Confocal stack maximum intensity projections of immunostainings for shTFs in the brain (top 3 rows) and VNC (bottom 4 rows). The brain midline is indicated with a vertical dotted line and the approximate contours of the optic lobes are highlighted with a dotted circumference in the top two rows. VNC top row: maximum intensity projections of confocal stacks. Some VNCs are missing the abdominal ganglion. White rectangles indicate the approximate areas shown in the bottom 3 rows. VNC bottom 3 rows: single slices showing individual shTFs expression (top 2 rows) and their overlap (bottom row). Examples of co-expressing cells are labelled with asterisks. Scale bars: brain and VNC 50µm, VNC insets 20µm.

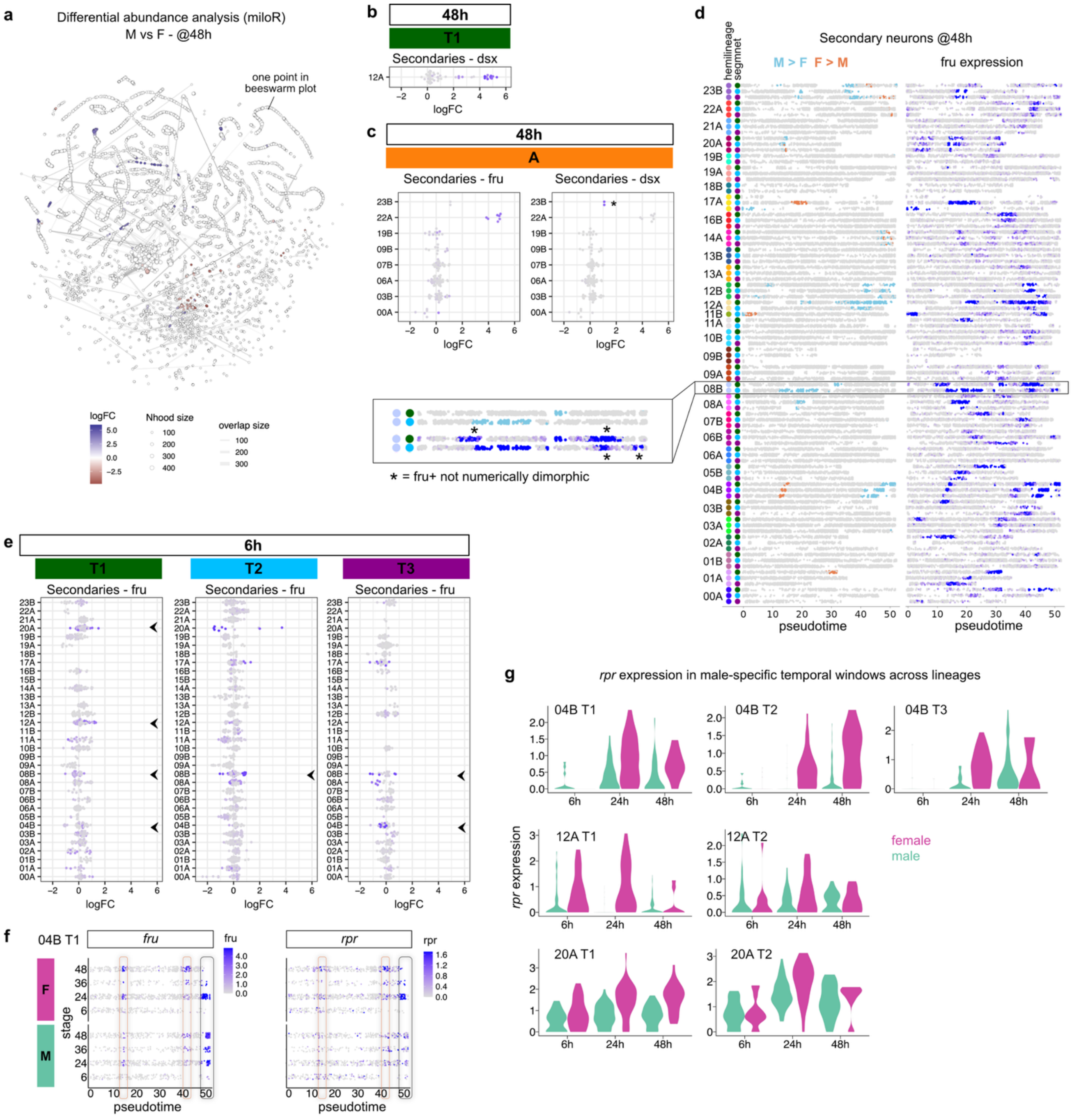


**Extended Data Fig. 12 | Development of sexual dimorphisms**

**a,** Differentially abundant neighborhoods at 48h resulting from miloR analysis. **b,** Beeswarm plot coloured by mean *dsx* expression, enriched in male-biased neighborhoods in 12A-T1 at 48h. **c,** Beeswarm plots of abdominal secondary hemilineages at 48h coloured by mean *fru* or *dsx* expression. **d,** Comparison between differentially abundant regions and *fru* expression. Atlas cells are distributed along pseudotime and separated according to hemilineage and segment. Inset shows 08B hemilineage in T1 and T2 at a higher magnification. Asterisks refer to fru-expressing regions that are not numerically dimorphic. **e,** Beeswarm plots of thoracic segments at 6h coloured by mean *fru* expression showing minimal differentially abundant neighborhoods. Arrowhead points to major fru+ neighborhoods statistically significant at 48h. **f,** Dot-plot showing expression of *fru* (left) and *rpr* (right) in 04B-T1 secondary neurons arranged along pseudotime and split according to sex and animal stage. Male-enriched temporal window encircled in grey, female-enriched windows in orange. Cell bodies position in Fig. 6h supports these predictions. **g,** Expression of *rpr* in male-enriched pseudotime windows across hemilineages.

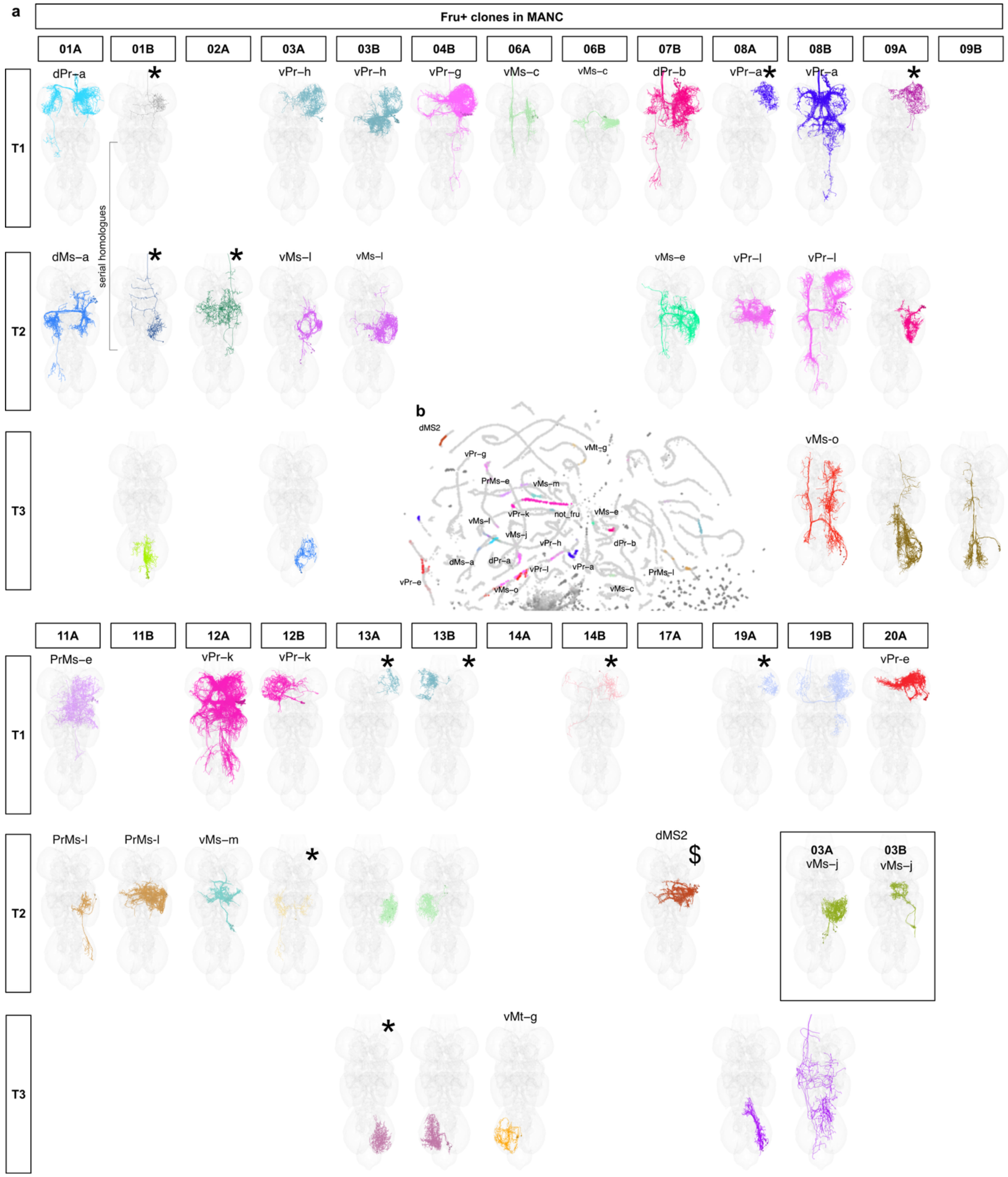


**Extended Data Fig. 13 | Fruitless secondary neurons in the atlas and the connectome**

**a,** Neurons belonging to fru clones in MANC connectome arranged by lineage and soma segment of origin. When the same clone includes cells from both hemilineages, the cells in the clone have been separated accordingly. Asterisks mark clones made by a single neuron. Dollar sign refers to *fru* secondary neurons previously annotated in the connectome, not identified as fru-clone. Clone name is reported for those cross-identified between connectome and transcriptome. Two different clones mapped onto 03 T2 lineage, potentially hitting cells born in different time windows. **b,** UMAP plot showing 48h atlas cells coloured by fru-clone identity.
